# CANE1-mediated autophagosome size regulation fine-tunes phosphate starvation tolerance in *Arabidopsis thaliana*

**DOI:** 10.64898/2026.02.12.705259

**Authors:** David Görg, Fiona Smith, Klara Machalett, Falko Koblenz, Ymy Thi Ngo, Sascha Höhne, Julian Arndt, Sylvestre Marillonnet, Nenad Grujic, Richard Imre, Karl Mechtler, Yasin Dagdas, Christin Naumann

**Affiliations:** Department of Molecular Signal Processing, Leibniz Institute of Plant Biochemistry, 06120 Halle (Saale), Germany; Institute of Biochemistry and Biotechnology, Martin Luther University Halle-Wittenberg, Halle (Saale), Germany; Department of Cell and Metabolic Biology, Leibniz Institute of Plant Biochemistry, 06120 Halle (Saale), Germany; Gregor Mendel Institute of Molecular Plant Biology, Vienna, Austria; Research Institute of Molecular Pathology (IMP), Vienna BioCenter (VBC), Vienna, Austria; Heidelberg University, Centre for Organismal Studies, Heidelberg, Germany

## Abstract

Inorganic phosphate (Pi) availability determines root development and plant performance. In *Arabidopsis thaliana*, external Pi is sensed by root tips and Pi limitation triggers ER stress–induced autophagy, yet the physiological impact and mechanisms controlling autophagosome biogenesis remain unclear. Here, we identify CANE1 (COMPONENT OF AUTOPHAGIC NETWORK) as a novel plant-specific regulator of autophagosome size in local Pi sensing. CANE1 associates with the plasma membrane-ER tethering protein VAP27-1 and binds to Pi-responsive ATG8 isoforms through conserved interaction motifs. Loss of CANE1 augments ER stress resistance by reducing autophagosome size in Pi-deprived root tips. Our findings establish CANE1 as a determinant of autophagosome size at the ER membrane. Thus, CANE1 functions as a molecular link between Pi-dependent ER stress signaling and fine-tuning of autophagic capacity.

## INTRODUCTION

Plant growth and development are tightly constrained by nutrient availability in soil. Phosphorus, primarily absorbed as inorganic phosphate (Pi), is the second most critical macronutrient, and Pi availability is often limited by complex chemical interactions with associated metals (e.g. Fe, Al) (Abel 2017; Abel and Naumann 2024; Lambers et al. 2015; Shen et al. 2011). To enhance Pi acquisition for maintaining cellular Pi homeostasis, plants undergo extensive developmental and metabolic reprogramming, which includes remodeling of root system architecture and chemical modification of the rhizosphere, respectively (Crombez et al. 2019; Giehl and von Wiren 2014; Kellermeier et al. 2014; Yang et al. 2024). In growing root tips of *Arabidopsis thaliana*, low external Pi rapidly reduces cell elongation and progressively inhibits cell division (Balzergue et al. 2017; Clua et al. 2024; Maniero et al. 2024; Muller et al. 2015; Naumann et al. 2022). The single ER-resident P5A-type ATPase PHOSPHATE DEFICIENCY RESPONSE 2 (PDR2) and the apoplastic ferroxidase LOW PHOSPHATE ROOT 1 (LPR1) form the core module of local Pi sensing in root tips (Svistoonoff et al. 2007; Ticconi et al. 2001; Ticconi et al. 2009). Decreasing Pi-availability activates LPR1 by facilitating access to its substrate (Fe^2+^), triggering cell-type-specific Fe^3+^ accumulation, formation of reactive oxygen species (ROS), and cell wall modifications, which interrupts cellular communication and consequently root growth (Balzergue et al. 2017; Gutierrez-Alanis et al. 2017; Muller et al. 2015; Naumann et al. 2022). PDR2 counteracts LPR1 function by maintaining root tip Fe homeostasis through an unknown mechanism (Jakobsen et al. 2005; Muller et al. 2015; Naumann et al. 2022; Naumann et al. 2019; Ticconi et al. 2009). Biochemical and structural studies reveal that P5A-type ATPases (P5As) act as membrane-helix dislocases that assist transmembrane protein insertion and remove mistargeted proteins, thereby safeguarding ER function and organelle integrity (Ji et al. 2024; Li et al. 2024; McKenna et al. 2022; McKenna et al. 2020). Disruption of P5A activity is detrimental across eukaryotes, affecting neuronal development in animals and causing pleiotropic ER-associated defects in yeast (Feng et al. 2020; Li et al. 2021; Qin et al. 2020; Sim and Park 2023). In plants, PDR2 function in root tips is linked to Pi-dependent ER stress-induced autophagy (Naumann et al. 2019; Stephani et al. 2020; Ticconi et al. 2009). Autophagy is a conserved degradation pathway essential for protein and organelle quality control, cellular homeostasis, and stress adaptation (Gross et al. 2025; Liu and Bassham 2012; Petersen et al. 2024; Su et al. 2020). Damaged organelles or protein aggregates are sequestered into double-membrane autophagosomes, typically 0.5–1.5 µm in diameter, which subsequently fuse with the vacuole for cargo degradation (Nakatogawa 2020; Zhao et al. 2022; Zheng et al. 2018). Autophagosome biogenesis originates at ER-associated phagophore initiation sites through PI3K complex (ATG6–VPS15–VPS34–ATG14) recruitment and progresses via ATG8 lipidation, mediated by the ATG12–ATG5–ATG16 complex. Additional regulators, such as SH3P2, interact with ATG8 and the PI3K complex to support phagophore expansion, cargo loading, vacuolar targeting, and possibly autophagosome size control (Nascimbeni et al. 2017; Walczak and Martens 2013; Zhuang et al. 2013). Selective autophagy receptors, including NBR1 (Jung et al. 2020; Svenning et al. 2011) and C53 (Stephani et al. 2020), deliver defined cargos to the phagophore through ATG8 interaction motifs (AIMs), and cargo characteristics can directly influence autophagosome dimension (Johansen and Lamark 2020). ATG8 isoforms enable organ- and stress-specific autophagy responses as illustrated by the differential complementation of carbon and nitrogen starvation phenotypes (Del Chiaro et al. 2025; Dong et al. 2025; Liu et al. 2021; Thompson et al. 2005). Beyond nutrient starvation, local Pi sensing in root tips and Pi deficiency in green tissues triggers iron-mediated ER stress-induced autophagy, likely originating from cargo overload within the secretory pathway (Naumann et al. 2019; Yoshitake et al. 2022). Notably, autophagosome size is highly variable and strongly shaped by the nature of the target, yet how autophagosome dimensions are controlled during biogenesis remains unknown. Emerging evidence suggests that ER-associated proteins at phagophore initiation sites are likely central to autophagosome size determination; however, the molecular mechanism and components involved remain elusive.

Here, using quantitative proteomics and interaction assays in Arabidopsis, we report a novel COMPONENT OF THE AUTOPHAGIC NETWORK (CANE1) as a plant-specific regulator of autophagosome size during Pi limitation. CANE1 associates with the ER–plasma membrane tethering protein VAP27-1 and the ER-membrane embedded glycosylation complex component DGL1, interacts with root tip-localized ATG8B and ATG8C isoforms via conserved AIM motifs, and its loss of function results in smaller autophagosomes and enhanced ER stress resistance. Through its role in Pi-mediated autophagy, CANE1 shapes autophagosome biogenesis, establishing an ER-associated regulatory network that converges with ATG8-mediated, organ-specific autophagy.

## RESULTS

### Phosphate limitation induces ATG8 isoform specific ER stress mediated autophagy

Tissue specific expression of ATG8 isoforms (ATG8A-ATG8I) in *Arabidopsis thaliana* mediate the autophagic response under distinct stress conditions (Del Chiaro et al. 2025; Dong et al. 2025). To investigate their roles in local root Pi sensing, we monitored native promoter-driven GFP-ATG8 protein fusions (*pATG8x::GFP-ATG8x*) (Del Chiaro et al. 2025) for autophagosome formation in wildtype (WT) seedlings transferred from Pi-replete (+Pi) to Pi-deprived (-Pi) conditions as well as after exposure to carbon starvation (-C) and tunicamycin-induced (+TM) ER stress (Fig. S1). Consistent with previous reports, GFP fluorescence was largely absent in basal (+Pi) condition for most ATG8 isoforms with the exception of ATG8A and ATG8F (Del Chiaro et al. 2025; Dong et al. 2025; Thompson et al. 2005). Upon carbon starvation, all tested lines exhibited GFP-positive puncta formation in the root transition zone and, to a lower extent, in the root tip, indicative of bulk autophagy activation independent of isoform specificity (Fig. S1). Likewise, TM treatment induced autophagosome formation in the transition zone of all transgenic GFP-ATG8 lines. Notably, Pi deprivation selectively activated autophagosome formation in root tips expressing GFP-ATG8 fusion isoforms B, C, and E suggesting an ATG8 isoform-specific involvement in Pi sensing (Fig. 1a, Fig. S1). Given that local root Pi sensing triggers ER stress-mediated autophagy in Pi limitation via the PDR2–LPR1 module (Naumann et al. 2019; Stephani et al. 2020), we introgressed *pATG8B/C::GFP-ATG8B/C* transgenes into the Pi-sensitive *pdr2* mutant and monitored autophagosome formation following Pi deprivation. When compared to +Pi conditions, the number of GFP-ATG8B labelled autophagosomes increased 2-fold and 2.5-fold in wildtype and *pdr2* roots, respectively, with no significant changes in *ATG8* transcript levels (Fig. 1, Fig. S2a). Interestingly, the number of GFP-ATG8C labelled puncta increased significantly only in *pdr2,* suggesting a specific role in ER quality control, which is supported by enhanced autophagosome formation upon TM treatment (Fig. 1). To assess ATG8 isoform-specific functionality, we examined local Pi sensing in the ATG8-deficient nonuple mutant *Δatg8* (Del Chiaro et al. 2025), alongside complementation lines expressing GFP-ATG8A or GFP-ATG8H, representing the two major ATG8 clades in *Arabidopsis* (Rogov et al. 2023). *Δatg8* mutants exhibited modest root growth defects in Pi deprivation, indicative of enhanced ER stress sensitivity (Fig. S2b). Notably, ectopic expression of either ATG8A or ATG8H did not rescue this phenotype, implying that autophagosome induction in local Pi sensing may involve isoform-specific functions.

**Figure 1:**
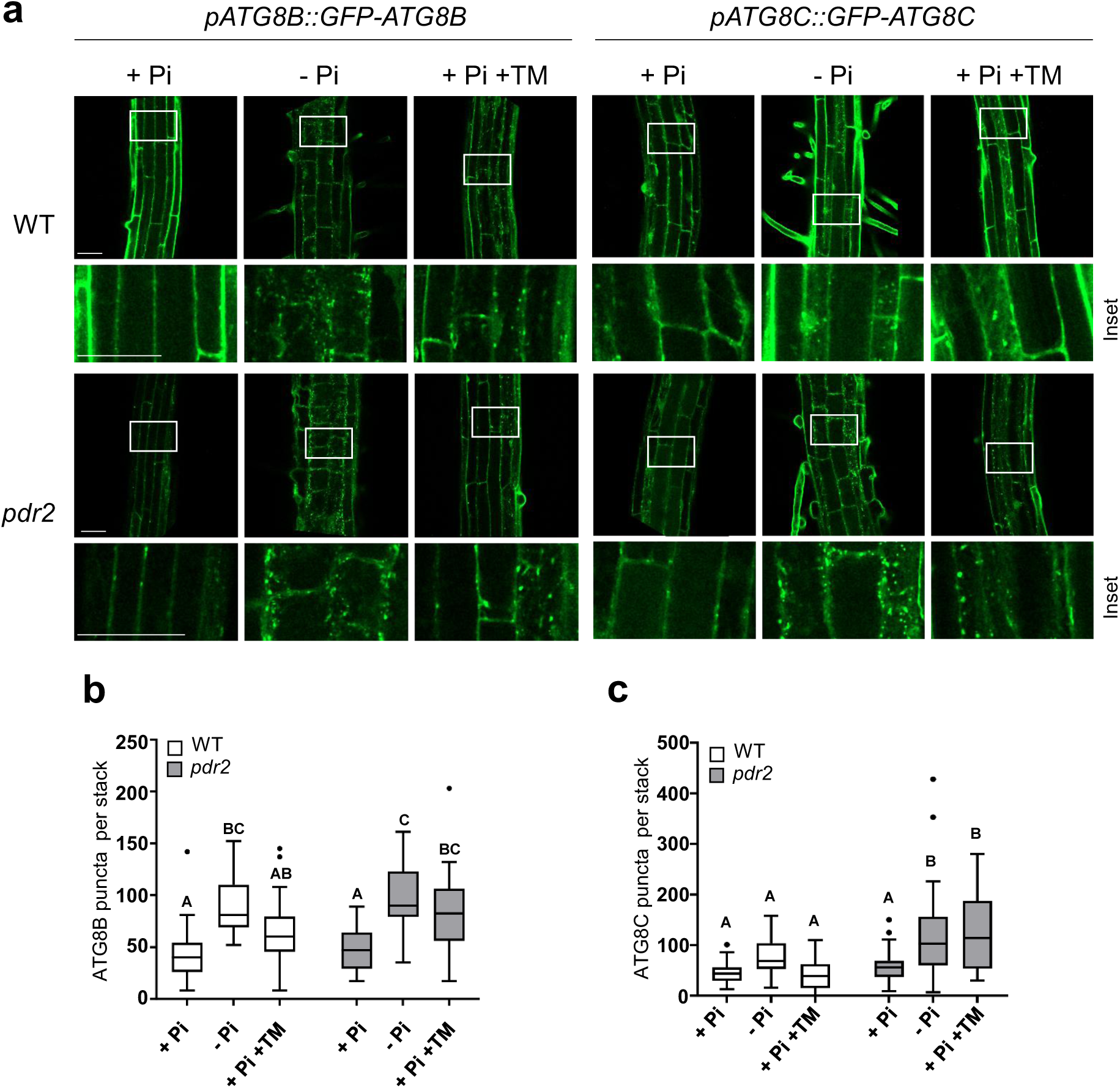
ER stress-induced autophagy by local Pi-sensing specifically activates ATG8B and ATG8C decorated autophagosomes. **a,** GFP-ATG8B and C labeled autophagosomes in Pi-deprived root tips. GFP-ATG8B/C derived fluorescence in primary root tips of transgenic (*pATG8B/C::GFP-ATG8B/C*) wildtype and *pdr2* seedlings after germination on +Pi agar (5 d) and subsequent transfer to –Pi medium for 0-24 h or treatment with 5 µg/mL tunicamycin (TM) for 6 h. Shown are representative images from three independent experiments fused Z-stacks of the transition zone. Rectangle indicate inset selection. Scale bars, 50 µm. **b** and **c**, Quantification of dataset shown in (**a**). Quantification of GFP-ATG8 puncta in root cells per normalized area. Box plots show medians and interquartile ranges of detected autophagosomes; outliers (>1.53 interquartile range) are shown as black dots. Letters denote statistical differences in condition at p < 0.05 (Two-way ANOVA and Tukey’s HSD post hoc test) and n ≥ 27.

### Local Pi sensing remodels autophagy-dependent root tip proteome

To elucidate the biological role of ER stress-induced autophagy in local Pi sensing and uncover novel components, we employed two complementary mass spectrometry-based screening approaches. Our working hypothesis posits that proteins regulated in an autophagy-dependent mode during Pi sensing should (i) exhibit altered abundances in autophagy-deficient mutants such as *atg2*, and (ii) physically interact with ATG8 isoforms. First, we performed proteomic profiling of wildtype, *pdr2*, and *atg2* root tips using tandem mass tag-based quantitative mass spectrometry (TMT-MS). Seedlings were germinated in Pi-sufficiency and subsequently transferred to control (+Pi) or ER stress-inducing (−Pi or +TM) conditions. Comparative analysis revealed that either ER stress condition caused pronounced proteomic shifts in root tips, with *atg2* displaying a significantly higher number of differentially abundant proteins relative to wildtype and *pdr2* (Fig. 2a, Fig. S3a, Data S1). We next analyzed the protein abundances in ER stress conditions in *atg2* in comparison to WT or *pdr2*. For Pi-deprivation 89 proteins were increased in *atg2* but not in wildtype or *pdr2* root tips, suggesting autophagy-dependent regulation (Fig.2b, Fig. S3b). This set includes core autophagy components such as ATG4, multiple ATG8 isoforms, and selective autophagy receptors, NBR1 and C53, previously implicated in ER stress responses and Pi signaling (Picchianti et al. 2023; Stephani et al. 2021; Stephani et al. 2020; Wild et al. 2014). Enrichment of these proteins reinforced the validity of our experimental design and highlights a functional role of autophagy in maintaining proteostasis during local Pi perception. Second, to identify Pi-responsive ATG8 interactors, we performed immunoprecipitation coupled with mass spectrometry (IP-MS) using wildtype and *pdr2* lines expressing *p35S::GFP-ATG8A* (Fig. S3d, e; Data S2). This screen revealed over 300 potential interactors, including ATG8B and ATG8C as well as NBR1 and C53, which are established ATG8 interactors (Stephani et al. 2020; Wilfling et al. 2020; Zhao et al. 2022). To discover functionally relevant ATG8-associated proteins in Pi-deprivation, we cross-referenced the ATG8A interactome (Fig. 2c,d) with the set of 89 proteins stabilized in *atg2* (Fig. 2b), which revealed ATG8B, ATG8C, NBR1 and an uncharacterized protein (AT5G40690), among the most enriched candidates across both datasets (Fig. 2e). Consistent detection of AT5G40690, suggests a novel function in autophagy-dependent Pi sensing and root proteostasis. We therefore refer to AT5G40690 as CANE1 (COMPONENT OF AUTOPHAGIC NETWORK).

**Figure 2:**
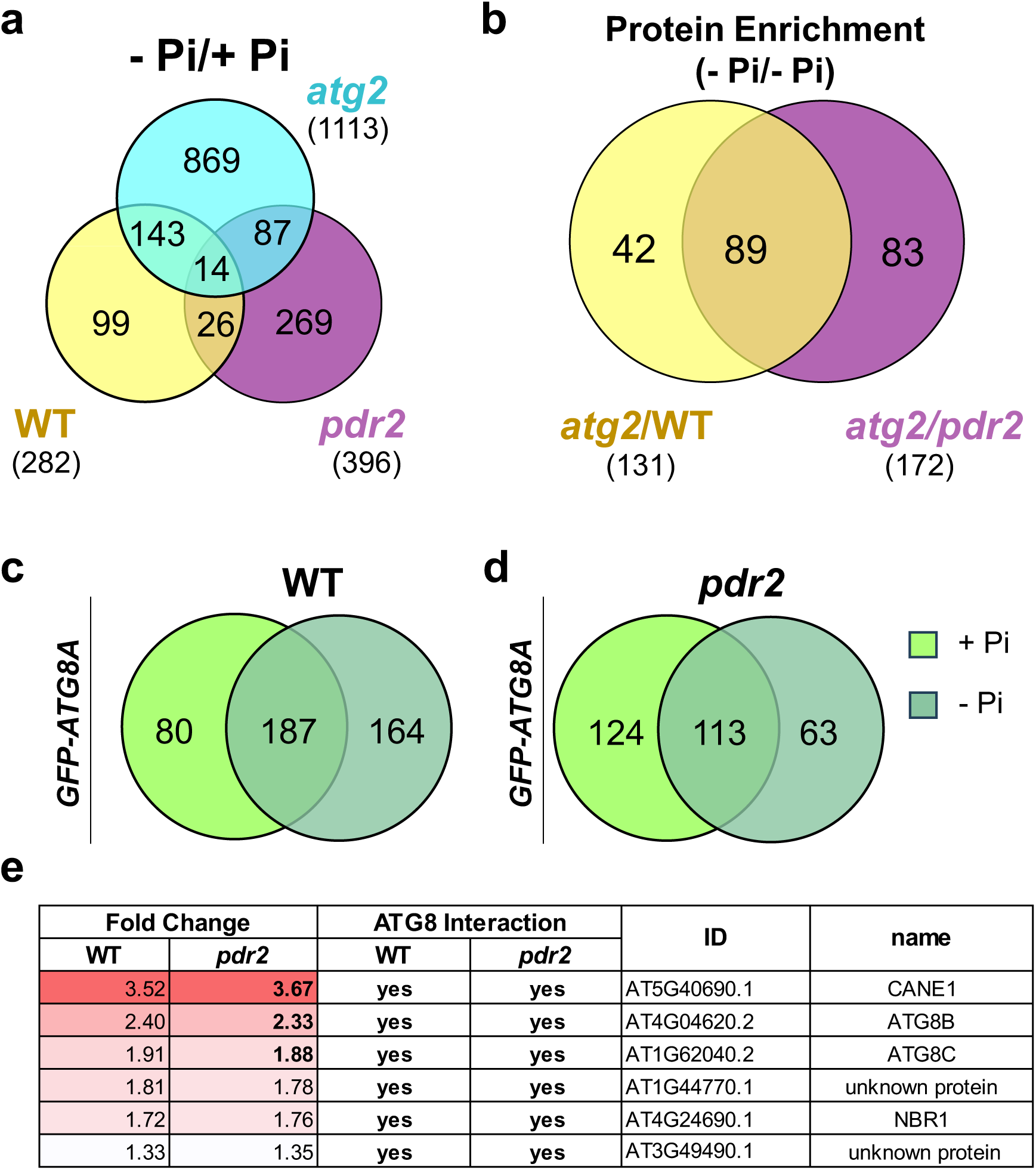
Autophagy modulates root tip proteome composition. **a,** Seedlings were germinated 5 days prior to transfer to the indicated treatments. 24 h post transfer primary root tips were excised and either directly subjected to extraction and quantification by mass-spectrometry (TMT-analysis) or used for co-immunoprecipitation followed by MS analysis. Venn diagram of three overlapping pairwise comparisons for root tip TMT-MS conducted in wildtype (yellow), *pdr2* (magenta) and *atg2* (cyan) plants: wildtype (-Pi) vs wildtype (+Pi) yellow cycle, *pdr2* (-Pi) vs. *pdr2* (+Pi) magenta cycle, *atg2* (-Pi) vs. *atg2* (+Pi) cyan cycle. Protein abundances with 0.6 > FC > 1.3. Total numbers are depicted in brackets. **b**, Overlap between proteins detected in –Pi conditions enriched in *atg2* compared to wildtype (yellow) or *pdr2-1* (magenta) identified by quantitative MS, FC > 1.3. Intersections show proteins with increased abundance in *atg2* but not in wildtype or *pdr2-1*. **c**,**d** *In vitro* interactome of GFP-ATG8A. Root extracts of GFP-ATG8A in transgenic wildtype (WT, **c**) and *pdr2* (**d**) primary roots were incubated with GFP-Trap beads and interacting proteins were analyzed by MS. Venn diagram of proteins co-eluting with GFP-ATG8A in +Pi (light green) and -Pi (dark green) according to the genotype. **e**, Summarized results of data obtained in c and d. Fold-Change (log2) of normalized protein abundances in *atg2* compared to wildtype (first column) and *pdr2* (second column) in -Pi conditions.

### CANE1 is evolutionally conserved among land plant species

While the core autophagy machinery is conserved across plants, the identification of modulators fine-tuning the autophagic pathway remains an active area of investigation (Cadena-Ramos and De-la-Pena 2024; Gross et al. 2025; Stephani and Dagdas 2020). AT5G40690 encodes a small, 210-amino-acid protein, expressed from a single exon. Phylogenetic analysis revealed that CANE1 is exclusive to and conserved across mostly all major land plant lineages apart from ferns and gymnosperms (Fig. 3a, Fig. S4). Notably, in a subset of angiosperms, including both monocot and dicot species but not bryophytes and *Amborella trichopoda*, we identified additional gene homologs encoding truncated proteins (∼110 amino acid residues), that lack the C-terminal domain present in canonical CANE1 orthologs (Fig. 3a, Fig. S4). In Arabidopsis, two such paralogs, HRG1 (AT2G411730) and HRG2 (AT5G24640) have been implicated in reactive oxygen species (ROS)-mediated root meristem maintenance (Gong et al. 2021). These findings suggest functional divergence or neofunctionalization of HRG proteins within specific angiosperm lineages such as the Brassicaceae.

**Figure 3:**
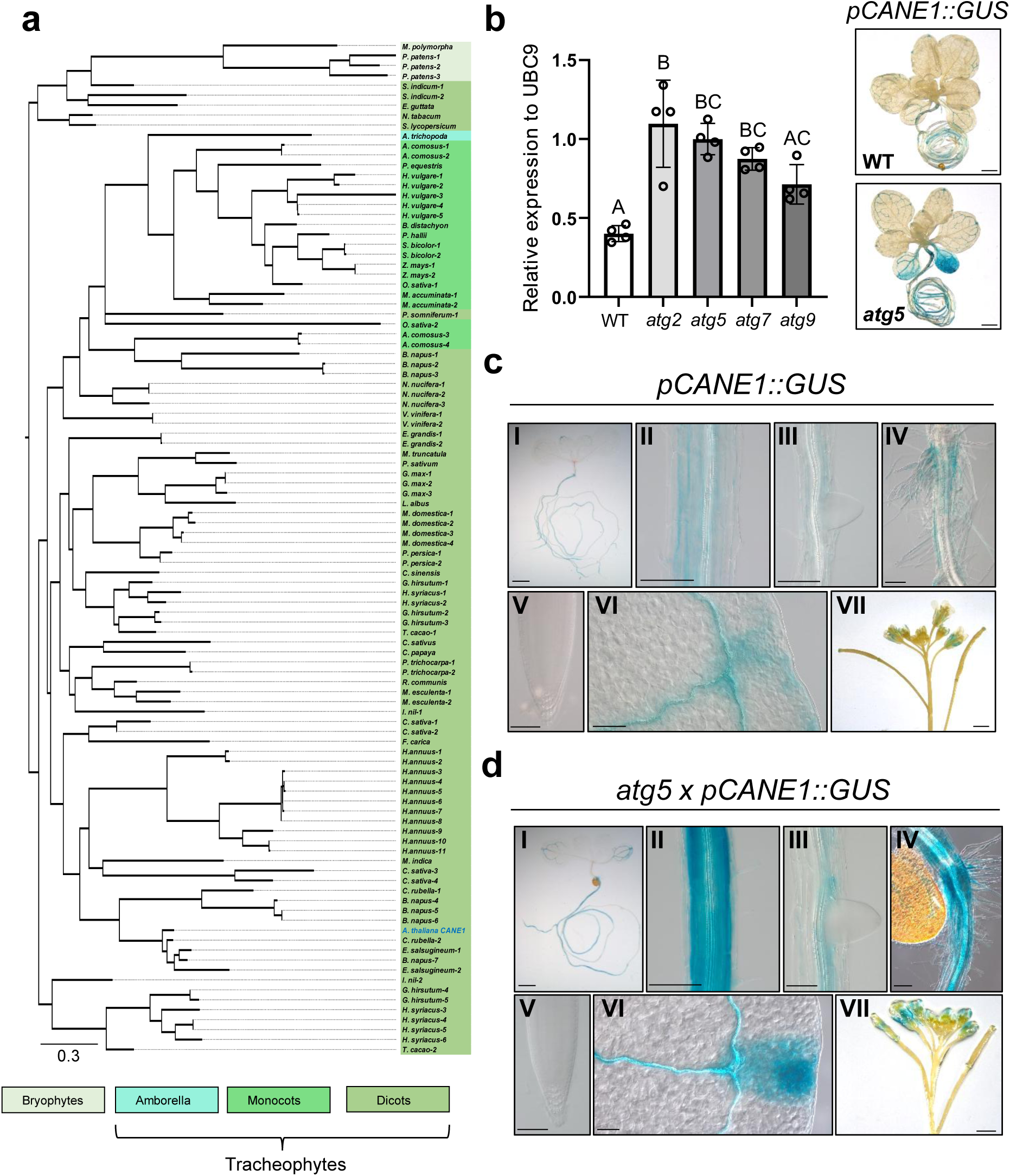
CANE1 is a plant specific protein. **a**, Maximum-likelihood midpoint-rooted tree (400 bootstrap replicates) of full-length CANE1 orthologues (see Supplementary Data 2). **b-d**, *CANE1* gene is predominantly expressed in root tissue and elevated in autophagy mutant plants. **b**, Left panel: RT-qPCR analysis of *CANE1* transcript abundance relative to *UBC9* in seedlings of wildtype, *atg2*, *atg5*, *atg7* and *atg9,* n = 4; right panel: *pCANE1::GUS* activity in 3 week old seedlings of wildtype and *atg5,* scale bar 1 cm. **c,d**, *pCANE1::GUS* activity in 7 days old wildtype (c) and *atg5* (d) seedlings (I), root elongation zone (II), lateral root emergence (III), root-shoot junction (IV), primary root tip (V), true leaf tip (VI) and flower (VII), I-VI scale bar 100 µm, VII scale bar 1 cm, n ≥ 8.

Expression analysis in whole seedlings revealed consistent upregulation of *CANE1* transcript levels in autophagy-deficient mutants (*atg2*, *atg5*, *atg7*, and *atg9*) compared to the wildtype control (Fig. 3b). To investigate organ- and tissue-specific *CANE1* expression, we generated transgenic *pCANE1::GUS* wildtype and *atg5* mutant lines (Fig. 3b-d). In wildtype, *CANE1* promoter activity was predominantly detected in root tissues, including lateral roots and root hairs, with increased activity in the root-shoot junction, while only weak GUS reporter signals were observed in the tip region of cotyledons and true leaves as well as in flowers and developing siliques (Fig. 3b,c). Interestingly, in the *atg5* background, the spatial distribution of *CANE1* promoter activity was comparable to wildtype, but markedly enhanced as evidenced by intense GUS staining, which correlates to elevated *CANE1* transcript levels measured in autophagy-deficient lines (Fig. 3b, d). Collectively, the evolutionary conservation of *CANE1*, its dominant expression in roots, and its transcriptional activation in autophagy mutants support a role for *CANE1* in nutrient-dependent ER stress responses in land plants.

### CANE1 interacts with ATG8 in an AIM-dependent manner

Co-immunoprecipitation of endogenous ATG8 with CANE1-mCherry expressed in Arabidopsis roots experiencing ER stress (Fig. 4a) validated our initial IP-MS identification of CANE1 as an ATG8 interactor (Fig. 2). We combined structural modeling with *in vitro* and *in vivo* studies to determine the ATG8-binding site of CANE1. AlphaFold2 modelling indicated CANE1 to be intrinsically disordered and predicted the presence of five putative ATG8-interacting motif (AIM) consensus sequences (W/F/V-X-X-L/I/V), with AIM5 (aa 202-205) being the best candidate followed by AIM4 (aa 183-189) (Fig. 4b; Fig. S5). Yeast two-hybrid interaction assays indicated AIM5-dependent CANE1 binding to all ATG8 isoforms, and strong interaction with ATG8G, H, and I (Fig. 4c). Transient expression of GFP fusions of CANE1 in tobacco (*Nicotiana benthamiana*) and live cell imaging showed predominantly cytosolic localization with partial net-like association reminiscent of ER structures (Fig. 4d). Importantly, the subcellular localization observed for GFP-CANE1 was not altered by introducing AIM4 and/or AIM5 deletions (Fig. 4e). Subsequent co-immunoprecipitation analysis demonstrated the dominant role of AIM5 in CANE1 for ATG8 binding. While GFP-CANE1 interacted with all RFP-ATG8 isoforms in tobacco leave extracts, which was also true for GFP-CANE1^ΔAIM4^, deletion of AIM5 (GFP-CANE1^ΔAIM5^) abrogated CANE1 interaction with most ATG8 isoforms except for ATG8E and ATG8G. As expected, deletion of AIM4 and AIM5 abolished CANE1^ΔAIM4/5^ binding to any ATG8 isoform (Fig. 4f; Fig. S6). Collectively, our results demonstrate differential interaction of CANE1 with ATG8 isoforms, which is primarily mediated by AIM5.

**Figure 4:**
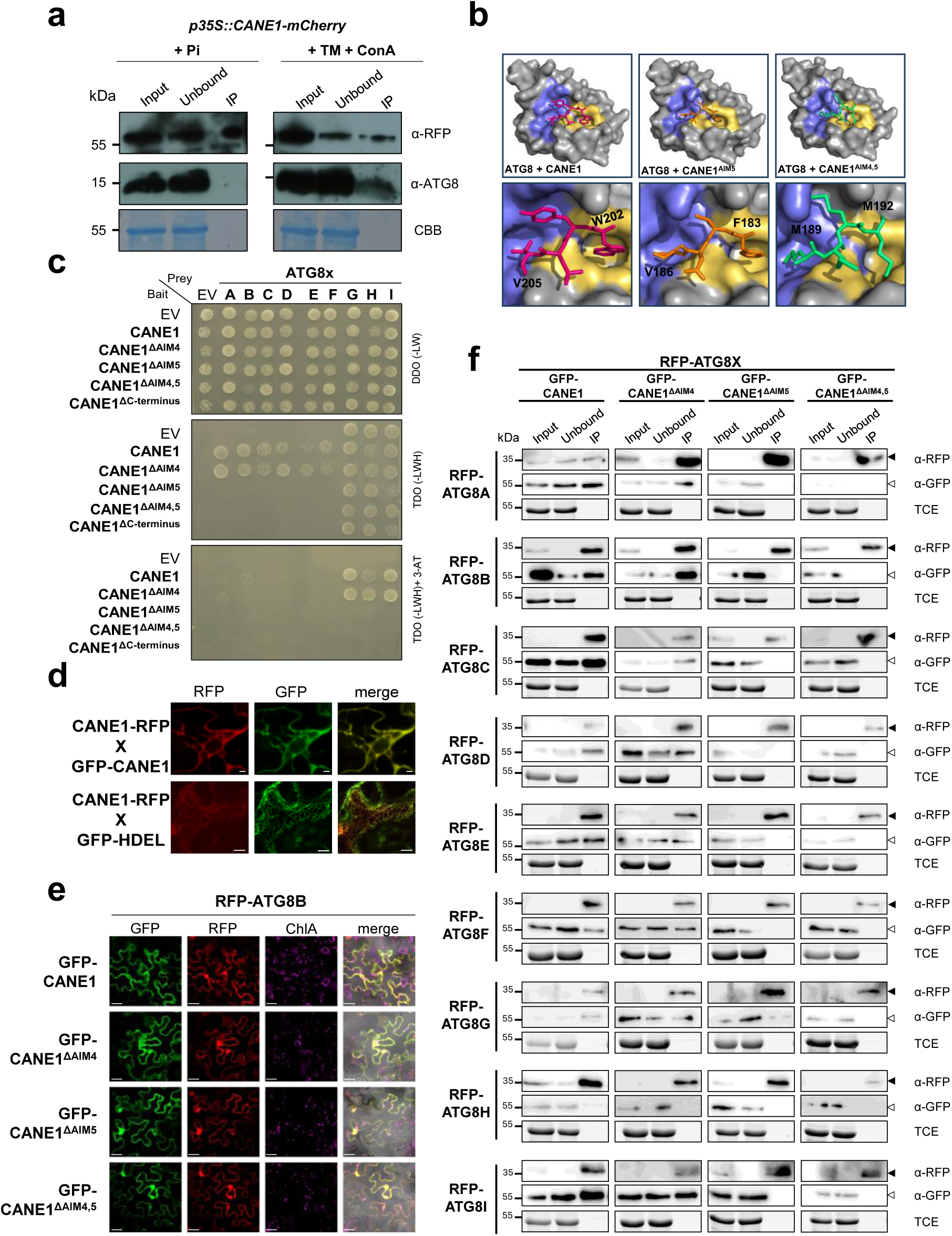
CANE1 interacts with ATG8 isoforms via select AIMs. **a,** endogenous ATG8 interacts with CANE1. *In vivo* immunoprecipitation of extracts of Arabidopsis seedlings expressing CANE1-mCherry after germination on + Pi media for 5 days. ER stress was applied by incubation in +Pi with 5 µg/mL TM and 1 µM ConA containing liquid media for 7h. Input and bound proteins were visualized by immunoblotting with anti-RFP and anti-ATG8 antibodies, loading control Coomassie blue staining (CBB). **b**, Colabfold prediction of CANE1-ATG8 interaction via AIM4 and AIM5. *In silico* prediction of CANE1 interaction with ATG8E revealed interaction via AIM5 (stick representation; see also Figure S6) which locates in Arabidopsis ATG8E (surface representation–grey) AIM pockets, known as W-site (site 1 –yellow) and L-site (site 2 –slate blue) (top panel) interacting with W202 and V205 (top right panel, close view – red). *In silico* mutation of W202A and V205A (CANE1^ΔAIM5^) predicts ATG8E CANE1 interaction via AIM4 (middle panel, right close view –orange) with F183 and V186 interacting with site 1 and site 2 respectively. CANE1 depleted of AIM4 and AIM5 is not predicted to interact with ATG8E via the AIM pockets (lower row panels, CANE1 ^ΔAIM4/5^ –green). **c**, CANE1 interacts AIM specific *in vivo*. Y2H assay between a GAL4 DNA binding domain (DBD, bait) fused variant of CANE1 with GAL4 activation domain (AD, prey) fused variants of all nine ATG8 isoforms A-I. GAL4-DBD and AD alone were included as negative controls (EV). The growth of yeast cultures on vector-selective (−Trp/−Leu) (top), interaction-selective (−Trp/−Leu/−His) (middle) and strong interaction-selective (−Trp/−Leu/−His + 3-aminotriazol) media (bottom) six days after spotting is shown. **d**, Subcellular localization of CANE1-RFP. Co-expression of CANE1-RFP with GFP-CANE1 (Upper row) or GFP-HDEL (lower row) in transient expression assays in *N. benthamiana*. The images are single optical sections of RFP (left column), GFP (center column) and merged signals (right column). Scale bars, 10 µm. **e**,**f** CANE1 interacts with ATG8 in an isoform and AIM specific manner. **e**, Localization of GFP-CANE1 variants (CANE1, CANE1^ΔAIM4^, CANE1^ΔAIM5^ CANE1^ΔAIM4/5^) transiently co-expressed with RFP-ATG8B in *N. benthamiana*. The images are single optical sections of GFP (first column), RFP (second column), chlorophyll A (third column, blue) and merged (including bright field) signals (last column). Scale bars, 30 µm, n = 3. **f**, *In vitro* co-immunoprecipitation assays of all nine RFP-ATG8x isoforms co-expressed with CANE1-GFP variants in *N. benthamiana* via RFP-Trap. Presence of RFP-ATG8A-I and GFP-CANE1 variants was visualized by immunoblotting with anti-RFP and anti-GFP antibodies, loading control TCE visualization, n = 5.

### CANE1 functions in ER stress-dependent autophagy during local Pi sensing

To investigate the relevance of CANE1–ATG8 interaction *in vivo* and its role for root responses to ER stress, we monitored autophagosome formation in transgenic lines expressing GFP-tagged CANE1 (*pCANE1::CANE1-GFP; pUBQ10::GFP-CANE1*) (Fig. 5a-d, Fig. S7). Consistent with transient expression in tobacco (Fig. 4d, e), CANE1-GFP exhibited a diffuse cytoplasmic distribution under control conditions, occasionally forming low-intensity puncta, which remained unchanged when expressed in the low Pi-sensitive *pdr2* mutant (Fig. 5a, Fig. S7a,c).

**Figure 5:**
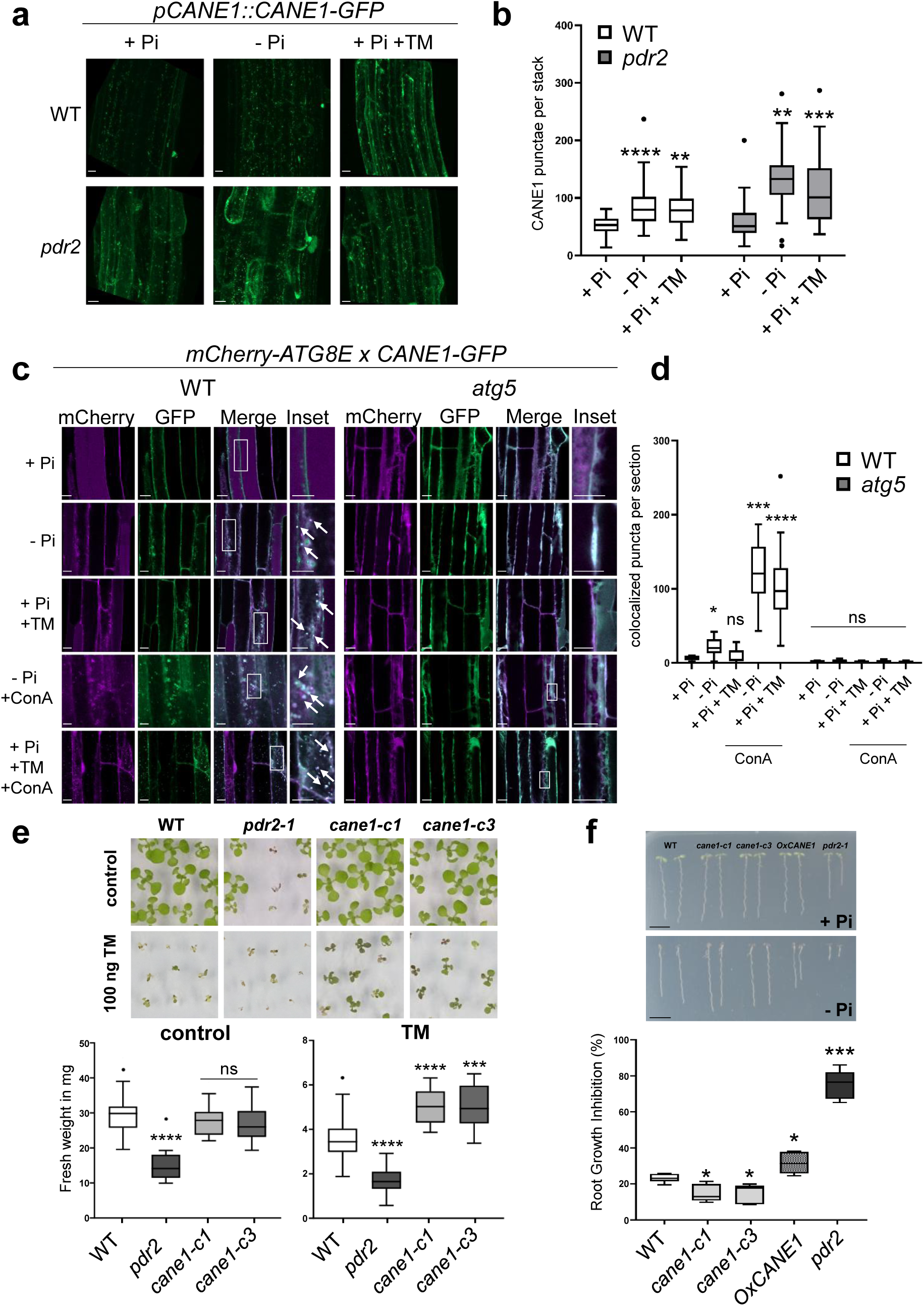
CANE1 functions in Pi-dependent ER stress-induced autophagy. **a,** CANE1-GFP derived fluorescence in the transition zone of transgenic (*pCANE1::CANE1-GFP*) wildtype and *pdr2* seedlings after germination on +Pi agar (5 d) and subsequent transfer to + or –Pi medium for 24 h or treatment with TM in liquid + Pi media for 6 h. Shown are representative images from three independent experiments fused Z-stacks of the root tip. Scale bars, 20 µm. **b**, Quantification of CANE1-GFP puncta in root cells per area. Box plots show medians and interquartile ranges of detected autophagosomes; outliers (>1.53 interquartile range) are shown as black dots, n ≥ 34. Significant differences compared to wildtype are indicated with * when p value ≤ 0.05, ** when p value ≤ 0.01, *** when p value ≤ 0.001, and **** when p value ≤ 0.001. Unpaired t-test with Welch’s correction. **c**, CANE1-GFP puncta colocalize with mCherry-ATG8E *in vivo*. Co-localization analyses of single plane confocal images obtained from transgenic wildtype (left panel) and *atg5* (right panel) Arabidopsis roots co-expressing mCherry-ATG8E (magenta) with CANE1-GFP (green). Seedlings were germinated 5 days prior transfer to the indicated treatment for 1 day (+/-Pi) or 5 h liquid media (+ Pi + TM; - Pi + ConA; + Pi + TM + ConA). Rectangles indicate Inset. Scale bars, 20 µm. **d**, Quantification of mCherry-ATG8E and CANE1-GFP colocalized puncta in the indicated treatments as described in a. n ≥ 5. **e-f**, loss of *CANE1* increases Arabidopsis ER stress tolerance. **e**, Top: 11 days old seedlings of wildtype, *pdr2*, *cane1-c1* and *cane1-c3* germinated on control or ER stress (100 ng/mL, TM) media. Bottom: Fresh weight of 5 pooled seedlings per sample, n = 17-18. **f**, Top: Germination of the indicated genotypes for eight days on Pi-sufficient (+Pi) or Pi-depleted (-Pi) conditions, scale bar = 1cm. Bottom: Primary root growth inhibition in percentage of the indicated genotypes, shown are summarized data of five independent biological experiments with each n ≥ 8. Significant differences compared to wildtype are indicated with * when p value ≤ 0.05, ** when p value ≤ 0.01, *** when p value ≤ 0.001, and **** when p value ≤ 0.001. Unpaired t-test with Welch’s correction.

Induction of ER stress by low Pi or TM treatment caused a rapid and robust increase in CANE1-GFP puncta and partial net-like localization with further enhancement in *pdr2* root cells (Fig. 5a,b). Importantly, CANE1 transcript levels and spatial expression in root tips remained unaffected in all lines tested (Fig. S7e,f). CANE1-GFP puncta co-localized with mCherry-ATG8E, demonstrating CANE1 association with autophagosomes (Fig. 5c,d, Fig. S7; (Feng et al. 2022)). Consistently, no CANE1-GFP puncta were detected in the autophagy-deficient *atg5* mutant although basal CANE1-GFP protein abundance was increased in *atg2* and *atg5* (Fig. S8d). These findings demonstrate that CANE1 associates with the ER and autophagosomes during ER stress in root tips, with a more pronounced response under Pi limitation compared to pharmacological ER stress induction by TM (Fig. 5a–d).

To investigate CANE1 function in *Arabidopsis*, we first analyzed T-DNA insertion (3’UTR) line (*SALK_092190*), which does not affect transcript abundance (Fig. S8a, b). We then generated two independent loss-of-function alleles using CRISPR–Cas9-mediated genome editing (Stuttmann et al. 2021): *cane1-c1*, carrying an insertion resulting in a frameshift and premature stop codon, and *cane1-c3*, harboring a deletion predicted to have a similar effect (Fig. S8b-c). Both mutant lines were phenotypically indistinguishable from wildtype during germination and early root development (Fig. 5e, f). Under ER stress conditions, however, both *cane1* loss-of-function alleles exhibited increased survival on TM-containing medium, indicating enhanced resistance (Fig. 5e). By contrast, no phenotypic differences were observed under carbon or nitrogen starvation, a condition commonly used to activate bulk autophagy (Fig. S8f-h).

We next examined the role of CANE1 in local Pi sensing. While both *cane1* mutants were significantly less sensitive to Pi deprivation, exhibiting enhanced primary root elongation, *CANE1* overexpression (*pUBQ10::GFP-CANE1*) displayed the opposite root growth phenotype on low Pi medium (Fig. 5f). However, the *CANE1* loss- and gain-of-function lines tested were indistinguishable with respect to iron and callose accumulation in root tips, two histochemical markers of local Pi sensing (Naumann et al. 2022) (Fig. S9).

Thus, our observations suggest that CANE1 is dispensable for bulk autophagy but functions in ER stress-induced autophagy modulating root growth response under local Pi limitation.

### CANE1 facilitates autophagosome biogenesis through ER tethering with VAP27-1

To elucidate a role of CANE1 in autophagy, we analysed the CANE1 interactome under basal and ER stress conditions. Under ER stress, CANE1 associates with ATG8C and factors involved in autophagosome maturation (ATG3, FREE1, SH3P2, VPS34, NBR1) and the ER membrane (VAP27-1, DGL1), implicating CANE1 function in phagophore expansion (Fig. 6a; Fig. S10a). Co-immunoprecipitation confirmed CANE1 interaction with NBR1 in wildtype and *atg5* roots, restricted to ER stress in wildtype and stabilized by ConA treatment (Fig. 6b). Considering the CANE1 localization and interactome under stress (Fig. 5c, Fig. 6a, Fig. S10a), we tested interactions of CANE1 with DGL1, an ER quality control component of the OST complex (Lerouxel et al. 2005), and with VAP27-1, an ER–PM tether involved in endocytosis and mitophagy (Li et al. 2022; Man et al. 2024; Stefano et al. 2018; Wang et al. 2019). CANE1 co-immunoprecipitated with DGL1 (Fig. S10b), supporting ER recruitment of CANE1. VAP27-1 also interacted with CANE1 in tobacco leaves and Arabidopsis roots under Pi limitation or TM-induced ER stress (Fig. 6c,d), suggesting that CANE1 may anchor phagophores to the ER via VAP27-1 interaction and, enable autophagosome membrane expansion. Notably, GFP-ATG8A puncta were smaller in *cane1-c1* root tips under Pi limitation, while NBR1 turnover and autophagosome amount remained unaffected (Fig. 6e,f, Fig. S10d, e), consistent with impaired phagophore expansion. CANE1 protein abundance remained unchanged in ER stress (Fig. S8e, S10c).

**Figure 6:**
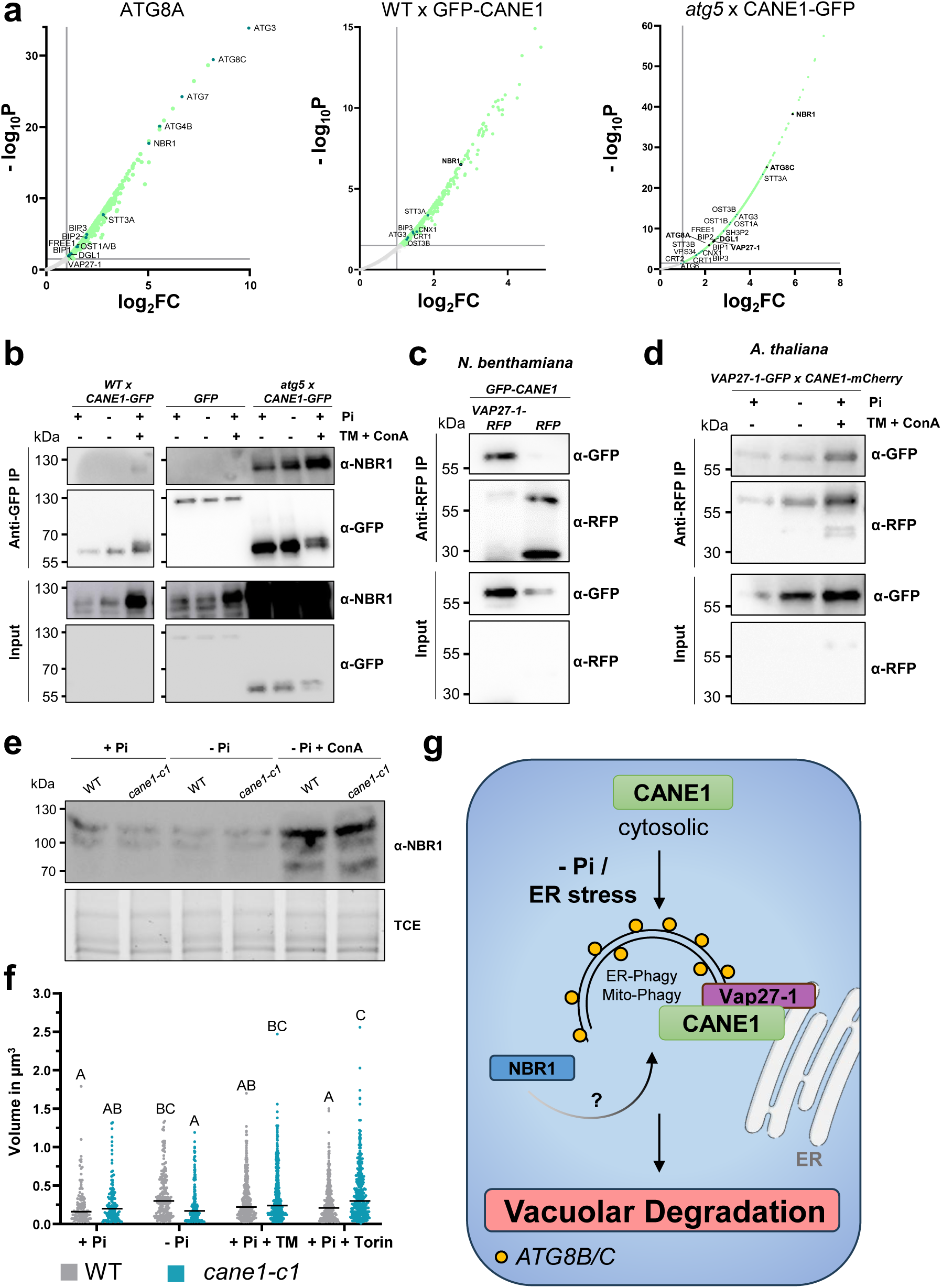
CANE1 interacts with NBR1 and VAP27-1 to modulate autophagosomal size. **a**, Enrichment of proteins interacting with ATG8A, *pUBQ10::GFP-CANE1* in wildtype and *pCANE1::CANE1-GFP* in *atg5* background in ER stress (+ TM + ConA) compared with a control expressing free GFP and represented by a volcano plot. Seedlings were germinated 10 days on liquid +Pi media prior to transfer to the indicated media. The horizontal grey line indicates the threshold above which proteins are enriched (P value <0.03, quasi-likelihood negative binomial generalized log-linear model), and the vertical dashed line indicates the threshold where the proteins’ log_2_FC is greater than 1. **b**, Western blot analysis from immunoprecipitation (IP) experiments using wildtype and *atg5* expressing *pCANE1::CANE1–GFP* or *pCANE1::GFP-GUS* to immunoprecipitate NBR1. Seedlings were germinated 10 days on liquid +Pi media prior to transfer to + or – Pi for 24 h or ER stress treatment with 5 µg/ml TM and 1 µM ConA for 6 h. Whole roots were harvested. The blots were probed with anti-NBR1 and anti-GFP antibodies to detect the presence of NBR1. **c-d**, CANE1 interacts with VAP27;1 in *N.benthamiana* and *A. thaliana* roots in ER stress. **c**, IP of *p35S::VAP27-1-RFP* or *p35S::mCherry-HDEL* co-expressed with *pUBQ10::GFP-CANE1* in *N.benthamiana* leaves. Input and bound proteins were visualized with anti-GFP and anti-RFP antibodies. **d**, IP of extracts of F1 Arabidopsis roots co-expressing *p35S::CANE1-mCherry* and *pVAP27-1::VAP27-1-GFP* after transfer for 24 h to Pi sufficient or deficient liquid media. ER stress was applied by incubation in +Pi with 5 µg/ml TM and 1 µM ConA containing liquid media for 6 h. Input and bound proteins were visualized by immunoblotting with anti-RFP and anti-GFP antibodies. **e**, NBR1 protein amount is not influenced by CANE1 in Pi-deprived roots. Endogenous NBR1 detection by anti-NBR1 antibody in extracts from wildtype and *cane1-c1* roots germinated 5 days on +Pi media prior to transfer to +Pi, –Pi or –Pi containing 1 µM ConA liquid media for 24h. **f**, Loss of CANE1 affects autophagosome size in Pi-stressed roots. Seedlings of wildtype or *cane1-c1* expressing *p35S::GFP-ATG8A* were germinated for 5 days on +Pi media prior to transfer for 24 h to +Pi or –Pi media or incubated with 5 µg/ml TM or 1µM torin for 6 h for autophagic induction. Size of GFP-ATG8A labelled puncta were measured via Arivis software and calculated volume was plotted accordingly. Scatter plot, median is shown as bold line, letters denote statistical differences at p < 0.05 (Two-way ANOVA and Tukey’s HSD post hoc test) and n = 20. **g**, Model of CANE1 function in autophagy. Induction of ER stress by local root tip Pi-sensing initiates CANE1 association to the ER and interaction with VAP27-1 to enable autophagosome maturation and membrane expansion for bulky cargo engulfment (e.g. ER-fractions, organelles). Potential NBR1 recruitment of CANE1 to the mature autophagosome by an unknown signal initiates ER-detachment and vacuolar degradation.

## DISCUSSION

Autophagy is essential for plant stress adaptation by maintaining cellular homeostasis under fluctuating environmental conditions, yet the mechanisms determining autophagosome maturation and size are largely unknown (Gross et al. 2025; Nakatogawa 2020; Zhao et al. 2022). In Arabidopsis, sensing of Pi limitation provokes iron-dependent ER stress response followed by selective autophagy induction in growing root tips and photosynthetic tissues; however the molecular determinants involved remain to be elucidated (Naumann et al. 2019; Yoshitake et al. 2021; Yoshitake et al. 2022). Here we identify CANE1 as a novel plant-specific regulator of autophagosome size during local root tip Pi-sensing. CANE1 associates with the ER through interaction with VAP27-1 and the OST complex subunit DGL1, which is essential for ER protein quality control (Lerouxel et al. 2005) (Fig. 6). Following ER stress perception, CANE1 associates with Pi-responsive ATG8 isoforms (B and C) preferentially via its C-terminal AIM motif (Fig. 1, 4). Loss of CANE1 leads to significant smaller autophagosomes and enhanced ER stress resistance (Fig. 5, 6) positioning CANE1 function in Pi-dependent ER stress signaling to the regulation of autophagosome biogenesis and maturation.

Phagophore expansion occurs at ER-associated initiation sites where lipid flux, membrane moulding, and cargo capture are coordinated (Gross et al. 2025; Liu et al. 2020; Liu and Bassham 2012; Nascimbeni et al. 2017; Su et al. 2020; Zhuang et al. 2013). The ER–PM tethering protein VAP27-1 is a key organizer of ER contact sites and supports mitophagy by promoting ER-mitochondria connections necessary for phagophore expansion around dysfunctional mitochondria (Li et al. 2022; Wang et al. 2019). VAP27-1 and its yeast homologues further support ER-stress adaptation through interactions with soluble ER-phagy receptors (Nthiga et al. 2020; Zhao et al. 2020). Although loss of VAP27-1 or DGL1 severely compromises plant development due to their essential roles in membrane tethering and N-glycosylation (Lerouxel et al. 2005; Wang et al. 2016), loss of CANE1 results instead in enhanced ER stress resistance, indicating a more specialized function. The interaction of CANE1 with VAP27-1 and DGL1 (Fig. 6) suggests that CANE1 contributes to stabilizing phagophore–ER contacts, a process central to the controlled expansion of the autophagosome membrane. We propose CANE1 modulates phagophore elongation by reinforcing ER–membrane interfaces via VAP27-1 and acts together with Pi-responsive ATG8B/C isoforms to establish a root cell-specific expansion regime for controlling autophagosome size. Such a CANE1–VAP27-1–ATG8B/C complex may further provide the spatial framework necessary for PI3K complex recruitment, thereby coordinating phagophore growth with cargo encapsulation.

In the physiological context of growing root tips, the degradation of stress-induced cargo must be balanced with the preservation of resources, especially under limited Pi availability. The smaller autophagosomes observed in *cane1* likely favour rapid turnover of small or diffusible cargos during early ER stress perception via the LPR1-PDR2 module, conferring enhanced resistance to low-Pi stress. Conversely, CANE1-dependent enlargement of autophagosomes in wildtype may facilitate more efficient sequestration of larger, bulky cargo, albeit at reduced throughput. This size-throughput trade-off aligns with the gradual response of Pi–Fe-linked ER stress induction, where rapid turnover may facilitate cell adaptation and sustain plant survival (Naumann et al. 2022; Naumann et al. 2019). CANE1 probably functions separate of established regulators of autophagosome maturation and fusion, such as SH3P2 and FREE1, whose loss is lethal, as well as autophagy inhibitors including COST1 and Rubicon-related factors (Bao et al., 2020; Gao et al., 2015; Matsunaga et al., 2009; Walczak & Martens, 2013; Zhuang et al., 2013). This reveals CANE1 function at a previously unrecognized regulatory layer dedicated specifically to autophagosome size determination.

We propose that in roots CANE1 relocates to ER contact sites upon low Pi stress, engaging ATG8B/C and VAP27-1 to stabilize PI3P complexes and regulate phagophore elongation. Following autophagosome closure, NBR1 may recruit CANE1 for turnover, positioning CANE1 as a new local regulator of autophagosome expansion (Fig. 6g). Our findings unravelled a complex regulatory network opening new avenues to dissect cargo selectivity and stress-responsive remodelling of endomembrane systems for plant environmental adaptation.

## METHODS

### Plant Materials

All *Arabidopsis thaliana* lines used in this study originate from accession Columbia (Col-0). Mutant lines *pdr2-1 (Ticconi et al. 2009)*, *lpr1 lpr2* (Svistoonoff et al. 2007), *ire1a ire 1b* (Naumann et al. 2019), *atg5-4 (Suzuki et al. 2005)*, *atg2-1* (Wang et al. 2011), *atg7-3* (Lai et al. 2011), *atg9-4* (Shin et al. 2014), *Δatg8 (Del Chiaro et al. 2025)* and transgenic lines *p35S::GFP-ATG8A* (Thompson et al. 2005), *pdr2-1 x p35S::GFP-ATG8A* (Naumann et al. 2019), *35Spro::SpER-GFP-HDEL, pdr2-1 x 35Spro::SpER-GFP-HDEL (Ticconi et al. 2009), pUBQ10::mCherry-ATG8E, atg5-1 x pUBQ10::mCherry-ATG8E* (Stephani et al. 2020), *Δatg8* x *pUBQ10::ATG8A*, *Δatg8* x *pUBQ10::ATG8H* (Del Chiaro et al. 2025) were previously described. T-DNA insertion line SALK_092190 was obtained from the European Arabidopsis Stock Center (NASC). The *cane1-c1* and *cane1-c3* knockout lines were produced by CRISPR-Cas9 technology (Stuttmann et al. 2021). Guide RNAs for target sequences (AAGCGTGAAGGCAAACAACAC and AACCCGGAAAGCTTCAAACC) were designed with CHOPCHOP (Labun et al. 2019) and cloned into pDGE332_M1 and pDGE334_M2E constructs (Ordon et al. 2017; Stuttmann et al. 2021) and assembled in pDGE347 for transformation. Double mutants were generated by genetic crossing and genotyped by PCR. Primers used for cloning and genotyping are listed in Supplementary table 1 and all vectors are listed in Supplementary table 2.

### Molecular Cloning

The *AtDGL1* coding sequence (CDS) was synthesized by Thermo Fisher Scientific with three point mutations introduced to eliminate internal BpiI recognition sites. The CDS was flanked by BpiI sites to enable Golden Gate cloning and subcloned into the MoClo vector pAGM1287, generating the level 0 *AtDGL1* module pNAU7. The *Arabidopsis thaliana* Ubiquitin 10 (Ubi10) promoter was amplified by PCR from plasmid pRedRoot (Limpens et al. 2004) and cloned as MoClo level 0 promoter modules pAGM11323, flanked by BpiI sites GGAG and AATG for use as a standard promoter module, and pAGM11335, flanked by BpiI sites GGAG and CCAT for assembly of transcriptional units containing N-terminal fusions. An internal BpiI site within the promoter sequence was removed during cloning by introducing a nucleotide substitution via the PCR primers. For N-terminal GFP fusion constructs, modules pAGM11335 (pUbi10), pICSL30006 (N-terminal GFP tag, (Engler et al. 2014) pNAU7 (*AtDGL1*), and pAGM16841 (tNOS, (Gantner et al. 2018) were assembled in a single BsaI Golden Gate reaction into pICH47742 (Weber et al. 2011), yielding pUbi10::GFP-AtDGL1::tNOS.

For C-terminal GFP fusion constructs, modules pAGM11323 (pUbi10), pICSL50004 (C-terminal GFP tag, (Engler et al. 2014) pNAU7 (*AtDGL1*), and pICH41421 (tNOS,) (Engler et al. 2014) were assembled in a single BsaI Golden Gate reaction into pICH47742, yielding pUbi10::AtDGL1-GFP::tNOS.

All other constructs were cloned via Gateway Technology.

### Plant Growth and Chemical Treatments

Seeds were surface-sterilized and germinated on 1% (w/v) Phyto-Agar (Duchefa) containing 2.5 mM KH_2_PO_4_, pH 5.6 (high Pi or +Pi medium) or no Pi supplement (low Pi or –Pi medium) as reported (Naumann et al. 2022). Agar media deficient in other nutrients (C, N, or Fe) or supplemented with ER stress-agents were prepared as described (Naumann et al. 2022; Naumann et al. 2019). The agar was routinely purified by repeated washing in deionized water and subsequent dialysis using DOWEX G-55 anion exchanger (Ticconi et al. 2009). For carbon starvation experiments on soil, 6 weeks-old plants grown under short-day photoperiod were transferred to darkness for 6 days followed by 7 days of recovery in short-day condition (Naumann et al. 2019; Qi et al. 2023; Thompson et al. 2005). Nutrient limitation in aseptic conditions was performed as follows: For carbon starvation, seedlings were transferred to carbon (sucrose) deficient media 5 days after germination and kept in the dark for 12 days followed by 7 days of recovery on +Pi media. For nitrogen starvation, 5 days-old seedlings (after germination on +Pi) were transferred to nitrogen deficient media for 7 days.

### Chemical Treatments

Tunicamycin (TM) and torin treatment were performed by transferring 5-day-old seedlings (germinated on +Pi) to liquid media supplemented with 5 µg/mL TM, 1 µM torin or DMSO (solvent) for 5-7 h. Concanamycin A (ConA) treatment was added to a final concentration of 1µM for the indicated time points (3-5 h).

### Histochemical staining

GUS (β-glucuronidase) staining was conducted as described (Naumann et al. 2022). Seedlings were incubated in GUS-staining solution (50 mM Na-phosphate, pH7.2; 0.5 mM K_3_Fe(CN)_6_; 0.5 mM K_4_Fe(CN)_6_; 2mM X-Gluc; 10mM EDTA; 0,1% TritonX) at 37°C and subsequently cleared using chloral hydrate solution (8:2:1 chloral hydrate:glycerol: ddH_2_O). Iron detection, based on Perls staining coupled to diaminobenzidine (DAB) intensification (Perls/DAB staining) was performed as previously described (Muller et al. 2015) with minor changes to the protocol. Plants were incubated for 10 min in 2% (v/v) HCl, 4% (w/v) K-ferrocyanide (Perls staining), or K-ferricyanide (Turnbull staining). For DAB intensification, plants were washed twice (ddH_2_O) and incubated (15 min) in methanol containing 10 mM Na-azide and 0.3% (v/v) H_2_O_2_. After washing with 100 mM Na-phosphate buffer (pH 7.4), plants were incubated for 3 min in the same buffer containing 0.025% (w/v) DAB (Sigma-Aldrich) and 0.005% (v/v) H_2_O_2_. The reaction was stopped by washing with 100 mM Na-phosphate buffer (pH 7.4) and optically clearing with chloral hydrate, 1 g/ml 15% (v/v) glycerol.

### Confocal Microscopy

Samples were analyzed using a multizoom stereomicroscope (Nikon AZ100) for overview images and a Zeiss AxioImager bright field microscope for detail images. Confocal microscopy was done on a Zeiss LSM900. Imaging was performed with a ×20 air, ×40 water or x63 water immersion objective using sequential acquisition. For each experiment, 7-10 individual samples were observed in three independent approaches. Images were processed using ImageJ (Schindelin et al. 2012) software. Total puncta numbers were counted using the MTB Cell Counter (Franke et al. 2015). Colocalization and Pearsons coefficient were calculated using the BIOP FiJi Plugin.

Callose was stained for 1 h with 0.1% (w/v) aniline blue (AppliChem) in 100 mM Na-phosphate buffer (pH 7.2) and carefully washed twice. Fluorescence was visualized using a Zeiss LSM 900 confocal laser-scanning microscope (excitation 405 nm, emission 498 nm) in 100 mM Na-phosphate buffer (pH 7.2) (Muller et al. 2015).

### Autophagosome size determination

Five-day old seedlings expressing *35S::GFP-ATG8A* in WT or *cane1-c1* were transferred to +Pi or -Pi for one day followed by liquid treatments to induce ER-stress by TM (5 µg/ml) or torin (1 µM) treatments for 6 h. Primary root transition zones were imaged and Arivis Vision 4D 3.6.2 (arivis AG) was used for analysis. A custom pipeline to measure autophagosome volume was set up using the Blob Finder with the following parameters: average size 2 µm, probability threshold 55%, split sensitivity 50%. Sphericity was set to > 0.65 and a segment feature filter was applied to ≥ 0.6 µm^2^.

### Transient expression in *N. benthamiana*

Leaves of 6-week-old tobacco (*Nicotiana benthamiana*) plants were co-infiltrated with *A. tumefaciens* harboring the respective plasmid. After 3 d, leaf discs were analyzed by confocal laser scanning microscopy (GFP: excitation 488 nm, emission 493-537 nm; mCherry: excitation 555 nm, emission 582-631 nm; chlorophyll autofluorescence: excitation 633 nm, emission, 669-722 nm).

### Yeast-two-hybrid assay

To perform direct Y2H test, the full length CANE1 as well as CANE1^F183A, V186A,^ CANE1^W202A, V205A^ and CANE1^F183A, V186A, W202A, V205A^ (CANE1^ΔAIM4^, CANE1^ΔAIM5^ and CANE1^ΔAIM4,5^ respectively), were cloned into the pDEST32 (Invitrogen) vector using gateway technology, to create protein fusions with the Gal4 DNA binding domain. All nine Arabidopsis ATG8 isoforms were cloned into the prey pDEST22 vector to obtain Gal4 activation fusions.

The bait and prey vectors to be tested for interaction of the fusion proteins were co-transformed into *S. cerevisiae* strain PJ69-4a using the standard LiOAc method (Gietz and Schiestl 2007). The yeast was spotted on plates with vector selective media DDO (double drop out media: SD/-Leu/-Trp) and were grown for 4-5 days at 30°C. Two independent colonies of each transformation were inoculated in 1x TE buffer and incubated at 30°C at 180 rpm overnight. On the next day, 10 μL of the colonies were spotted on DDO, interaction selective media (triple drop out media: SD/-Leu/-Trp/-His) and triple drop out media containing 2 mM 3-aminotriazole (triple drop out media: SD/-Leu/-Trp/-His/+3-aminotriazole). Pictures were taken 6 days after spotting. SD plates were prepared with 1.6% (w/v) Bactoagar (AppliChem) and the SD media was obtained from TakaraBio.

### Affinity-based Immunoprecipitation

Proteins were extracted from plant material with RIPA-buffer (50 mM Tris-Cl, pH 7.6, 150 mM NaCl, 20 mM NaF, 10 mM Na_4_P_2_O_7_, 1 mM EDTA, 0.5 mM EGTA, 1% Nonidet-P-40, 0.5% Na-deoxycholate, 1 mM PMSF, 1 mM DTT) for 30 min at 4°C. Post centrifugation the cleared lysate was diluted 1:1 with detergent-free RIPA and incubated with equilibrated GFP-trap or RFP-trap Agarose beads (ChromoTek) for 2h at 8°C on a multi-rotator. After centrifugation (2500xg, 4°C) and washing 3-times (detergent-free RIPA), proteins were eluted at 68°C for 15 min in 2x SDS sample buffer. The supernatant was transferred to a new tube and boiled at 95°C for 5min.

### Immunoblotting

Seedlings were germinated on +Pi medium for 5 d prior to transfer to Pi-sufficient (+Pi) or deficient (-Pi) media. Whole roots were harvested 1dpt after treatments. For basal protein quantification whole seedlings were directly harvested 5dpg. Proteins were extracted using RIPA-buffer (50 mM Tris-Cl, pH 7.6, 150 mM NaCl, 20 mM NaF, 10 mM Na_4_P_2_O_7_, 1 mM EDTA, 0.5 mM EGTA, 1% Nonidet-P-40, 0.5% Na-deoxycholate, 1 mM PMSF, 1 mM DTT) for 30 min at 4°C. Protein concentrations were measured using the Pierce™ 660 nm Protein-Assay-Kit (Thermo Fisher, 22662). Equal amounts of total proteins were loaded on 10% SDS-gels containing TCE (2,2,2-Trichlorethanol, Sigma-Aldrich, T54801) as loading control (Ladner et al. 2004). All gels were simultaneously immunoblotted. All antibodies used in this study are listed in Supplementary table 3.

### Quantitative proteomics

Primary root tips were excised 24 h after transfer to the indicated media and collected into liquid nitrogen. Tissue lysis, sample preparation, protein labeling, Tandem-Mass-Tag (TMT) spectrometry, MS/MS data analysis, and TMT-quantifications were performed as recently described (Naumann et al. 2022; Rodriguez et al. 2020; Stephani et al. 2020).

NanoLC-MS/MS Analysis: The nano HPLC system (Vanquish *Neo* UHPLC-System, Thermo Scientific) was coupled to an Orbitrap Astral mass spectrometer, equipped with a Nanospray Flex ion source (all parts Thermo Scientific). Peptides were loaded onto a trap column (PepMap Acclaim C18, 5 mm × 300 μm ID, 5 μm particle size, 100 Å pore size, Thermo Scientific) at a flow rate of 25 μl/min using 0.1% TFA as mobile phase.

After 5 minutes, the trap column was switched in line with the analytical column (Aurora Ultimate C18 25cm × 75 μm ID, 1.7 μm particles, 120 Å, with integrated emitter, Ionopticks operated at 50°C). Peptides were eluted using a flow rate of 300 nl/min, starting with the mobile phases 98% A (0.1% formic acid in water) and 2% B (80% acetonitrile, 0.1% formic acid) and linearly increasing to 35% B over the next 60 minutes, followed by an increase to 95% B in 1.7 min, a 4-min hold at 95% B, and re-equilibration with 2% B for three column volumes (equilibration factor of 3.0).

The Orbitrap Astral was operated in data-dependent mode, performing a full scan in the Orbitrap every 0.7 s (m/z range 375–1500 m/z; resolution 240,000; standard AGC target, maximum injection time 10 ms, minimum intensity 5,000). MS/MS spectra were acquired in the Astral analyser by isolating 1.6 Da windows and fragmenting precursor ions with normalized HCD collision energy of 30% with a maximum injection time of 5 or 10 ms correspondingly or until a standard AGC target was reached. Fragment ions ranging from 150–2000 m/z were acquired. Precursor ions selected for fragmentation (include charge state 2-6) were put on a dynamic exclusion list for 30 s.

### qRT PCR

Abundance of mRNAs was quantified using fluorescence-based real-time RT-PCR and gene-specific amplimers listed in Supplementary table 1. Relative mRNA abundances were normalized relative to *UBC9* transcript abundance. Samples from three independent experiments were measured.

### Phylogenetic Analysis

CANE1 sequence alignment and phylogenetic analysis were done using BLAST search of curated sequences in the NCBI database and phytozome 13. Obtained sequences were filtered for fragments. All phylogenetic trees were calculated by sequence alignment using MAFFT 7108 with default settings and created at the CIPRES web-portal with RAxML 8.2.10109 for maximum likelihood analyses using the JTT PAM matrix for amino acid substitutions in RAxML. All related sequences used for phylogenetic analyses are provided (Supplementary Data 2).

### Quantification and Statistical Analysis

All claims of statistical significance can be found in the corresponding figure legends. Statistical tests were carried out using GraphPad Prism 10 for Windows.

## Acknowledgements

We thank P.J. Hussey and P. Wang for seeds; M. Ried and S. Abel for scientific discussions and critical reading of the manuscript. This work was supported by the Deutsche Forschungsgemeinschaft (DFG, German Research Foundation) projects 492493685 and 400681449/GRK 2498.; an EMBO Short-Term Fellowship to C.N. and institutional core funding (Leibniz Association) from the Federal Republic of Germany and the State of Saxony-Anhalt. Work in the K.M. group was financially supported by the EPIC-XS (project number 823839), the Horizon 2020 Program of the European Union, and the ERA-CAPS I 3686 project of the Austrian Science Fund.

## Author contributions

D.G. and C.N. designed research; D.G., F.S., K.M., J.A., F.K., S.M., Y.TN., N.G. and S.H. conducted experiments and analyzed data; K.M. and R.I. performed proteomic analysis; Y.D. and C.N. supervised the project; D.G. drafted the manuscript; C.N. wrote the article.

---

**Supplementary Figure S1:**
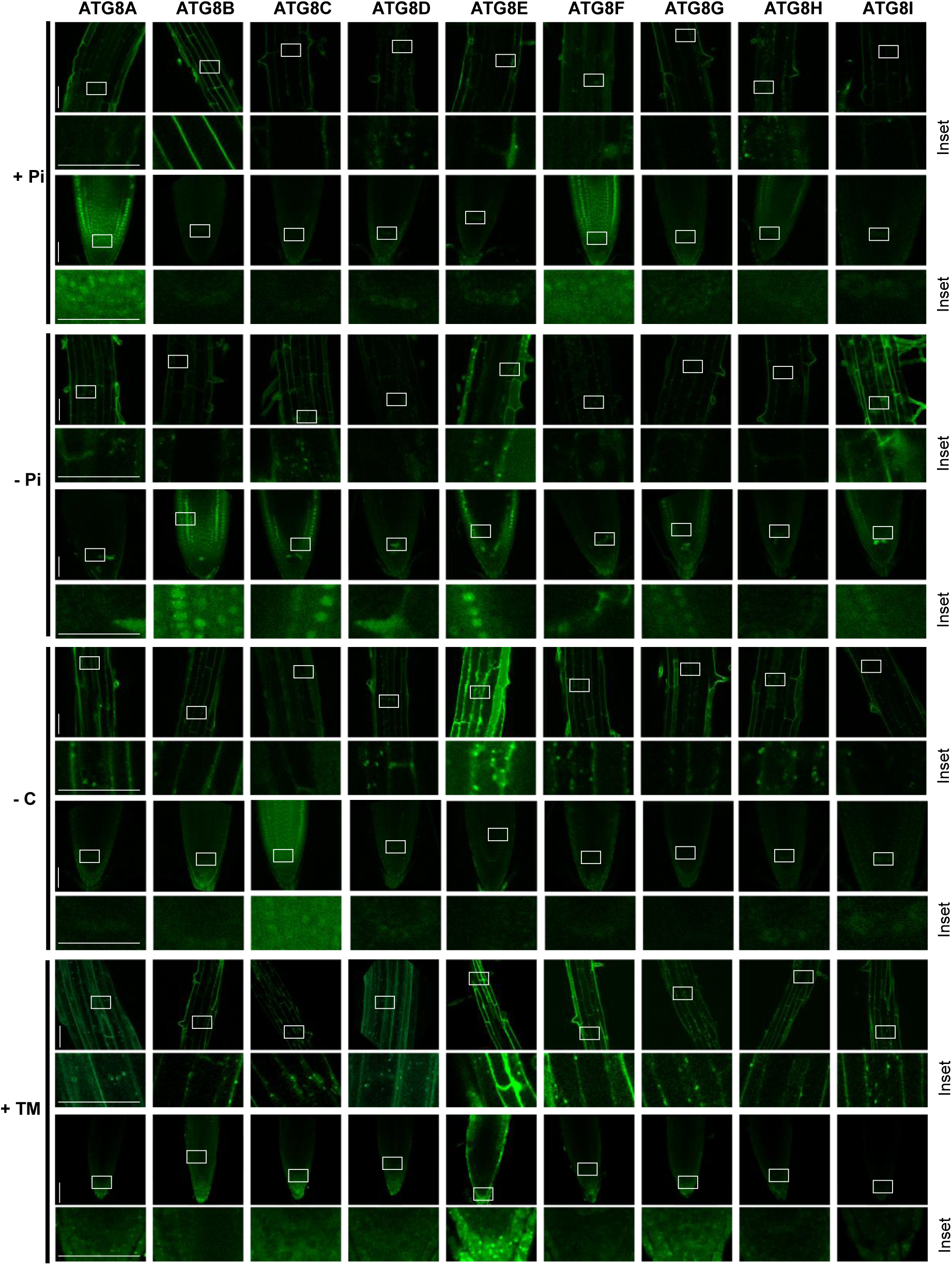
Root tip ATG8 isoform specific autophagosome decoration. Seedlings were germinated 5 days prior transfer to the indicated treatment. Representative confocal microscopic images showing autophagosome formation of all 9 ATG8 isoforms (*pATG8x::GFP-ATG8x*) in the transition zone (row 1,3,5,7) or primary root tip (row 2,4,6,8) challenged with local Pi deprivation (row 3 and 4), carbon starvation (row 5 and 6) for 24 h or 5 µg/mL tunicamycin (TM) treatment (row 7 and 8) for 6-8 h, rectangle indicate Inset selection, n ≥ 10.

**Supplementary Figure S2:**
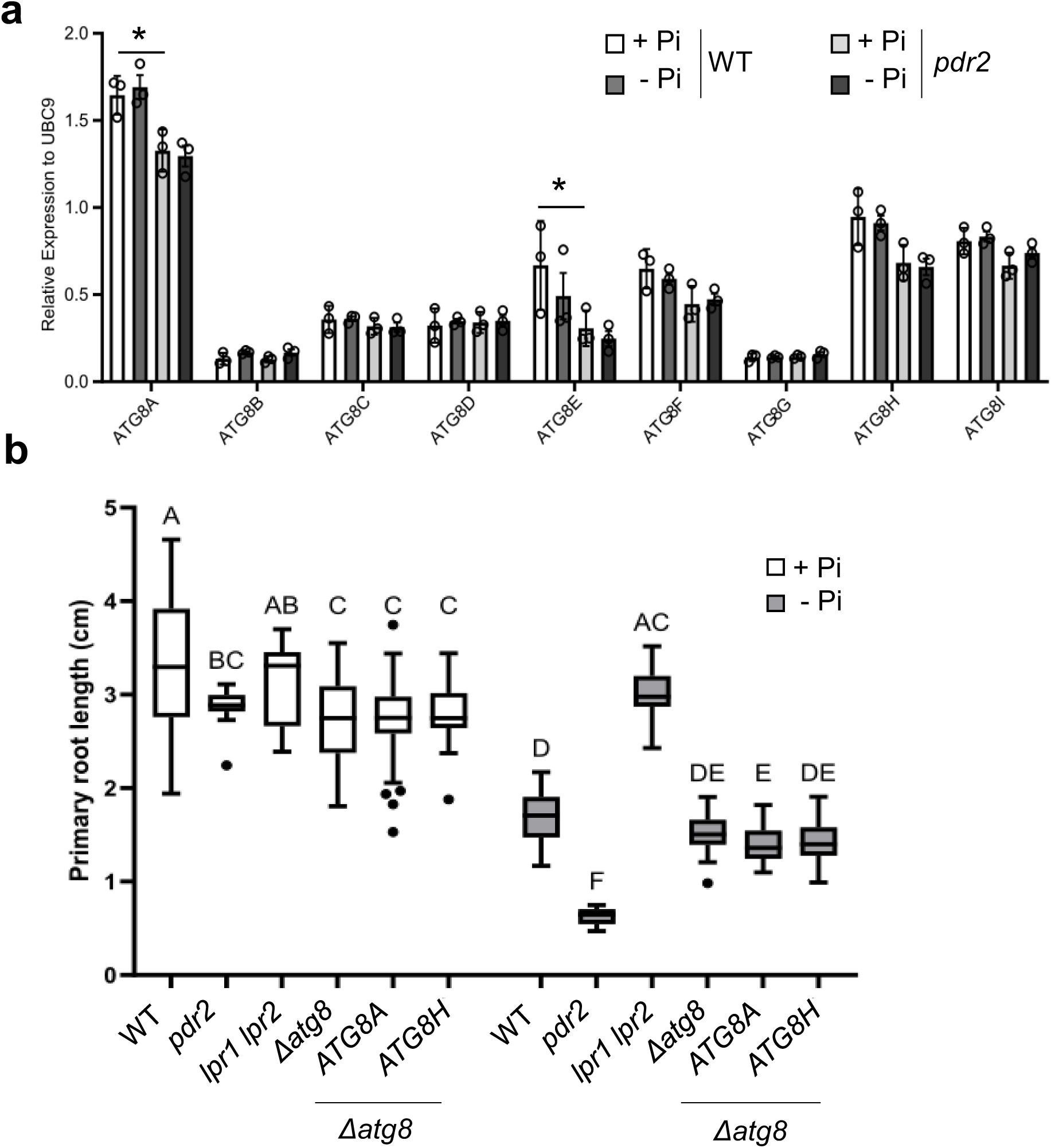
Select ATG8 isoforms are involved in root tip Pi-dependent ER stress-mediated autophagy. **a**, Analysis of *ATG8A-I* relative transcript levels (normalized to UBC9 expression) in wildtype and *pdr2* root tips after transfer to +Pi or –Pi medium for 24 h (germinated for 5 days on +Pi) ±SD; n = 3; Two-way ANOVA, Post-hoc Tuckey, *p value ≤ 0.05. **b**, ATG8A and ATG8H fail to complement *Δatg8* sensitivity to Pi limitation. Seedlings were germinated for 5 days on +Pi media prior to transfer to medium supplemented with or without Pi (-Pi, +Pi) and imaged after 4 days. Gain of primary root extension was measured. Box plots show medians and interquartile ranges of primary root gain; outliers (>1.53 interquartile range) are shown as black dots. Letters denote statistical differences in condition at p < 0.05 (Two-way ANOVA and Tukey’s HSD post hoc test) and n ≥ 13-50.

**Supplementary Figure S3:**
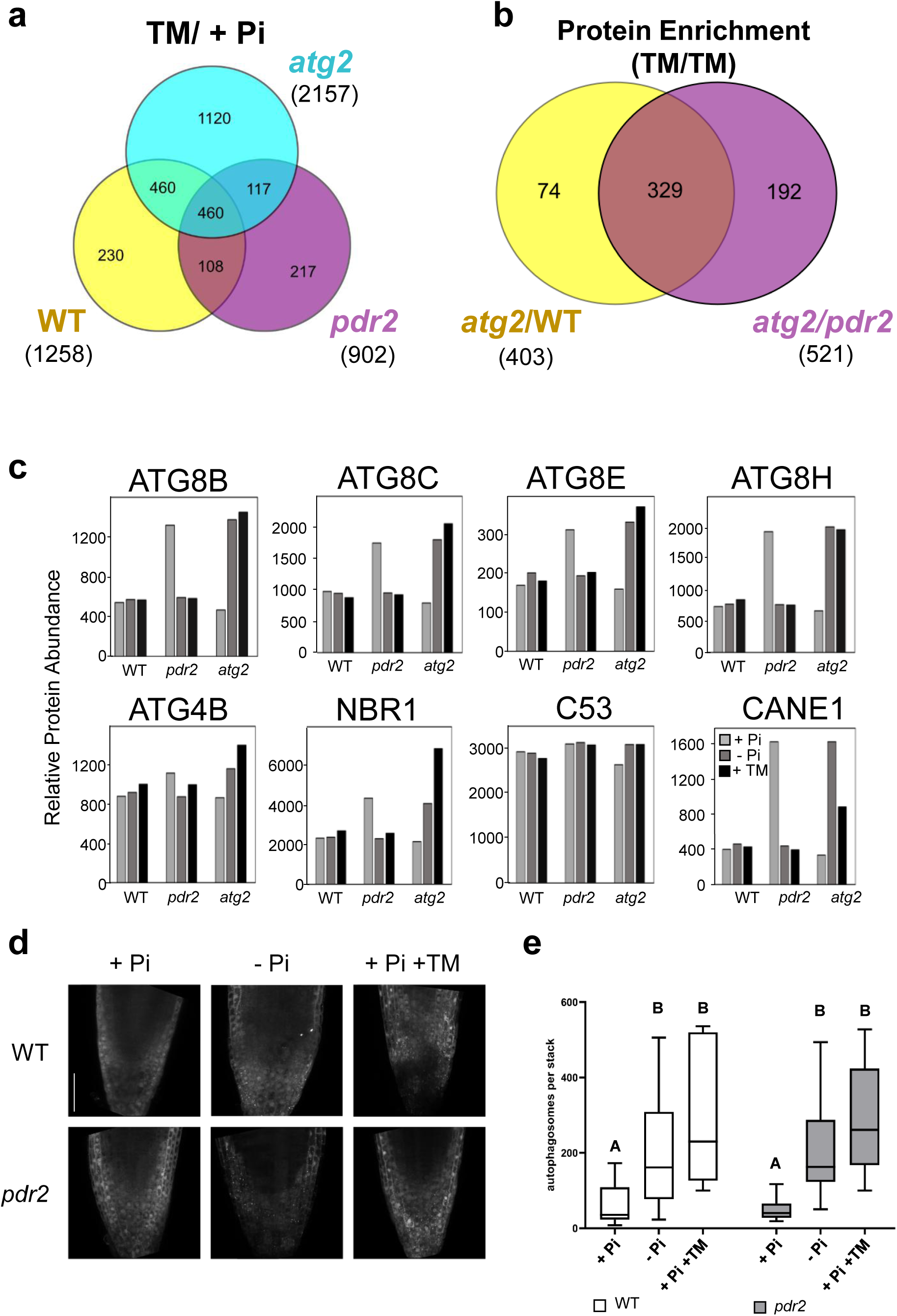
ER-stress derived adaptation of root tip proteomes. **a,** Chemically induced ER stress impacts root proteome. Venn diagram of three overlapping pairwise comparisons for root tip TMT-MS conducted in wildtype (yellow), *pdr2* (magenta) and *atg2* (cyan) plants: wildtype (+Pi) vs wildtype (+Pi+TM) yellow cycle, *pdr2* (+Pi) vs. *pdr2* (+Pi+TM) magenta cycle, *atg2* (+Pi) vs. *atg2* (+Pi+TM) cyan cycle. Protein abundances with 0.6 > FC > 1.3. Total numbers are depicted in brackets. **b**, Overlap between proteins identified in + Pi + TM conditions enriched in *atg2* compared to wildtype (yellow) or *pdr2-1* (magenta) identified by quantitative MS, FC > 1.3. **c**, Proteins of the autophagic pathway are affected in local Pi-mediated root tip proteomic adaptation. Plotted protein abundances of ATG8B/C/E/H, ATG4B, NBR1, C53 and CANE1 in +Pi (light grey), -Pi (dark grey) and +Pi +TM (black) conditions measured via TMT-MS in wildtype, *pdr2* and *atg2.* **d,e** Autophagosome induction in Pi deprived and ER stressed root tips. **d**, GFP-ATG8A derived fluorescence in primary root tips of transgenic (*p35S::GFP-ATG8A*) wildtype and *pdr2* seedlings after germination on +Pi agar (5 d) and subsequent transfer to –Pi medium for 24 h or treatment with 5 µg/mL TM for 6 h. Shown are representative images from three independent experiments fused Z-stacks of the root tip. Scale bars, 50 µm. **e**, Quantification of GFP-ATG8 puncta in root cells per normalized area and number of slices. Box plots show medians and interquartile ranges of detected autophagosomes; outliers (>1.53 interquartile range) are shown as black dots. Letters denote statistical differences in condition at p < 0.05 (Two-way ANOVA and Tukey’s HSD post hoc test) and n ≥ 6.

**Supplementary Figure S4:**
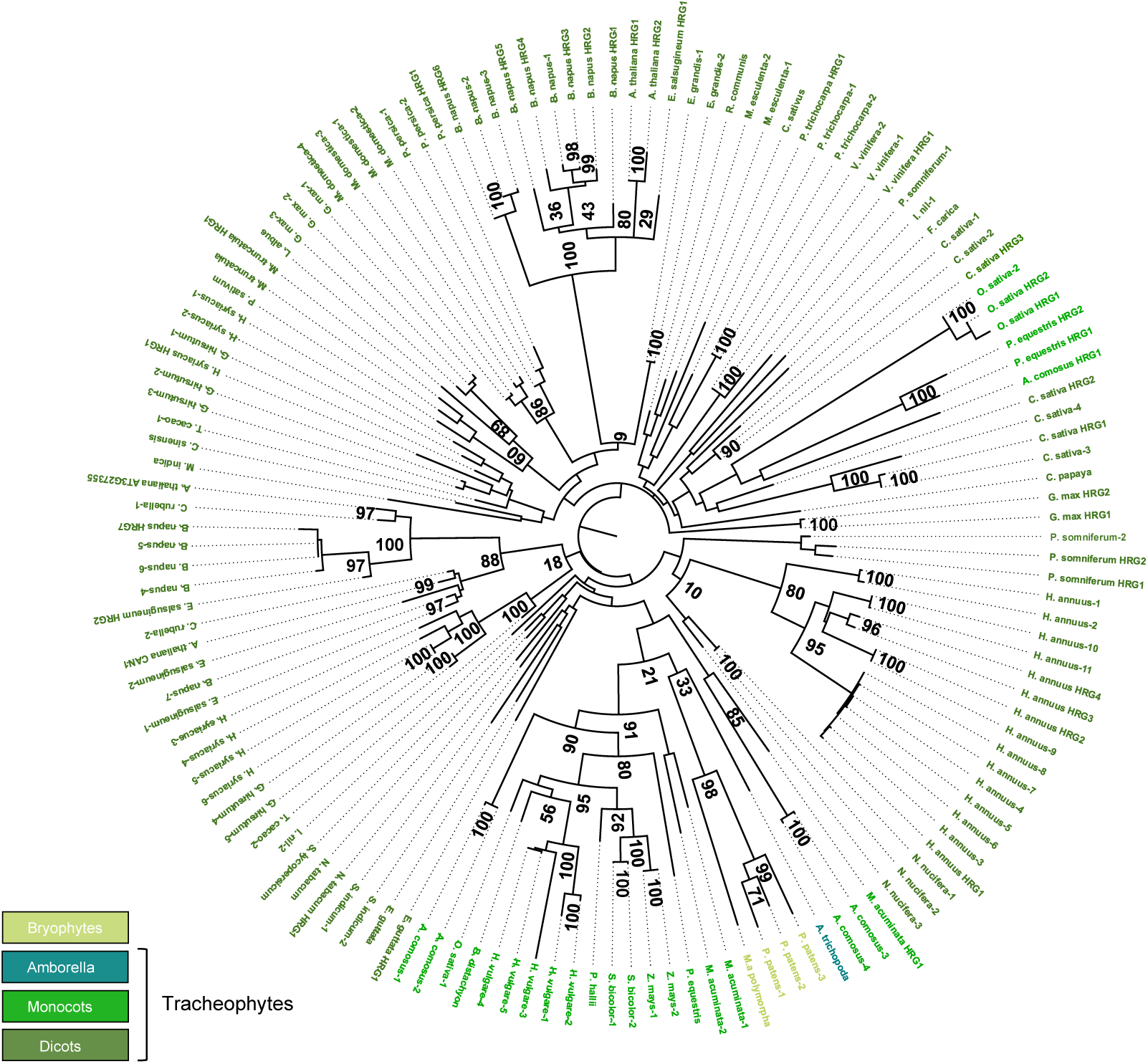
CANE1 is present in most land plant species. Maximum-likelihood midpoint-rooted phylogenetic tree (250 bootstraps) of 139 predicted CANE1-like proteins (Phytozome). Identified CANE1 like proteins of bryophytes (light green), amborella (turquoise), monocots (leaf green) and dicots (dark green). Full-length CANE1-like sequences are annotated by organism in a numeric order, C-terminally truncated proteins are annotated according to *A. thaliana* HRG1 and 2 respectively.

**Supplementary Figure S5:**
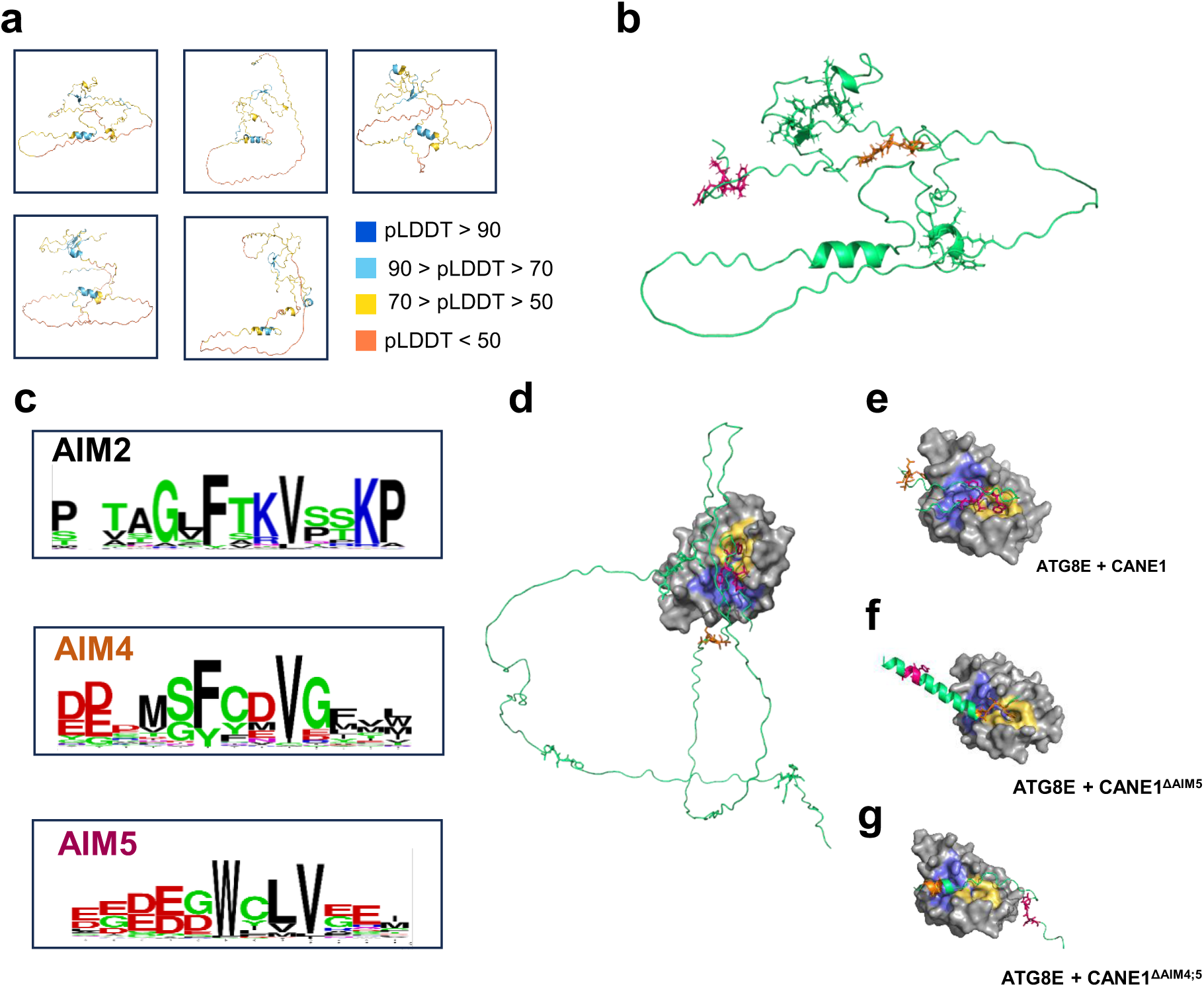
CANE1 is an intrinsically disordered protein. Structure and interaction prediction of CANE1 by AlphaFold2. **a**, CANE1 is intrinsically disordered. Top five structure prediction by AlphaFold2 colored according to the pLDDT score of the predicted structure. **b**, CANE1 structure in green cartoon, with all five identified AIMs in stick representation (AIM1-3 green, AIM4 orange, AIM5 magenta). **c**, Specific conserved AIMs, with AIM4 and AIM5 exclusively present in full-length CANE1 proteins. **d-g** Predicted AIM-dependent interaction of full length CANE1 with ATG8E. **d**, CANE1 (cartoon, AA 192-210) interacts with ATG8E (surface representation) via AIM5 (stick, magenta), **e**, ATG8E AIM pockets, known as W-site (site 1 –yellow) and L-site (site 2 –slate blue) interacting with W202 and V205. **f**, Predicted interaction of CANE1^ΔAIM5^ (cartoon, AA 181-210) via AIM4 (stick, orange) and **g**, loss of predicted CANE1 ATG8E interaction in CANE1^ΔAIM4,5^ (cartoon, AA 181-210).

**Supplementary Figure S6:**
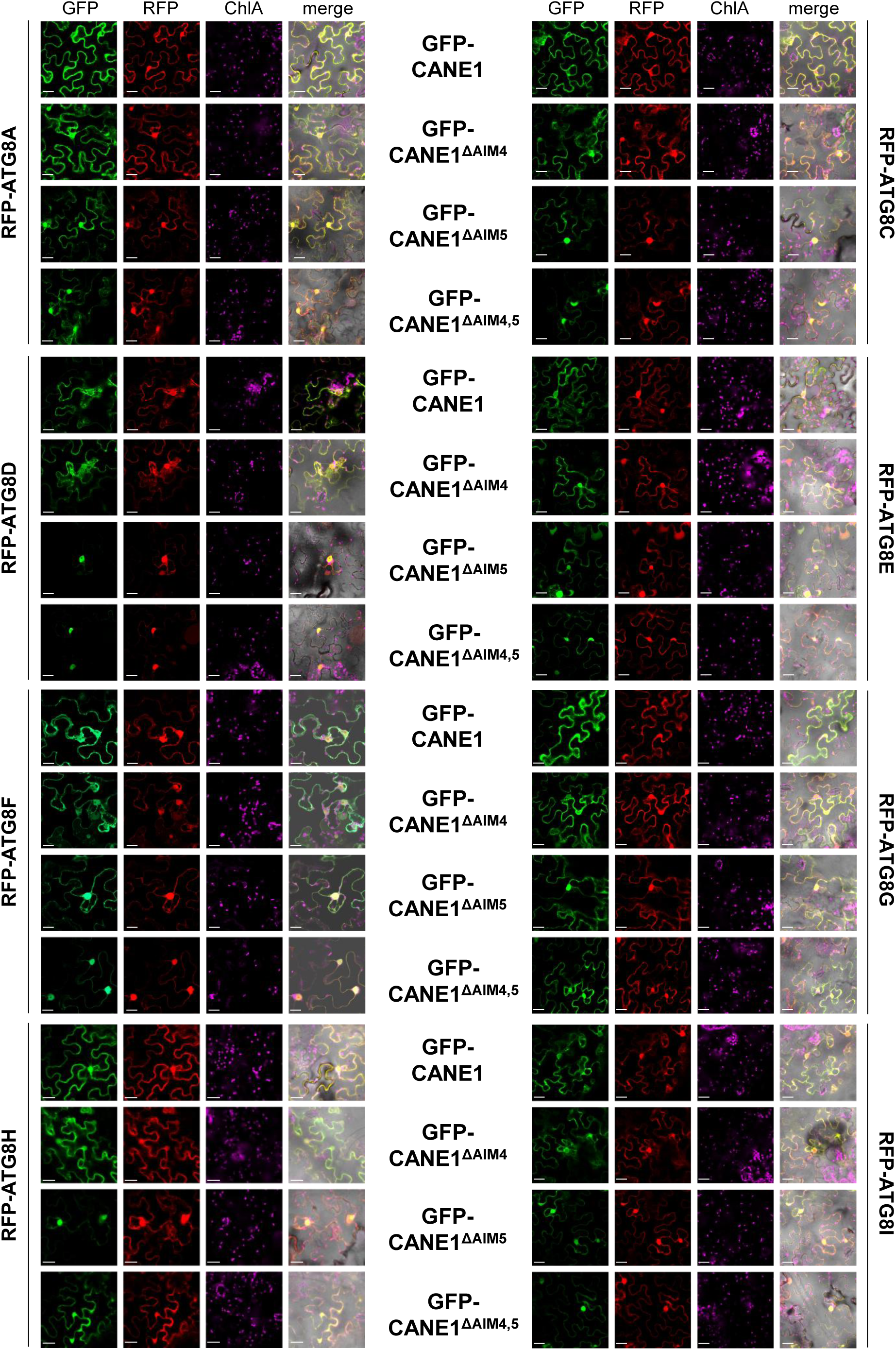
CANE1 co-localizes with all ATG8 isoforms in N. *benthamiana*. Localization of GFP-CANE1 variants (CANE1, CANE1^ΔAIM4^, CANE1^ΔAIM5^ CANE1^ΔAIM4/5^) transiently co-expressed with RFP-ATG8A, C-I in *N. benthamiana*. The images are single optical sections of GFP (first column), RFP (second column), chlorophyll A (third column, blue) and merged (including bright field) signals (last column). Scale bars, 10 µm; n ≥ 3.

**Supplementary Figure S7:**
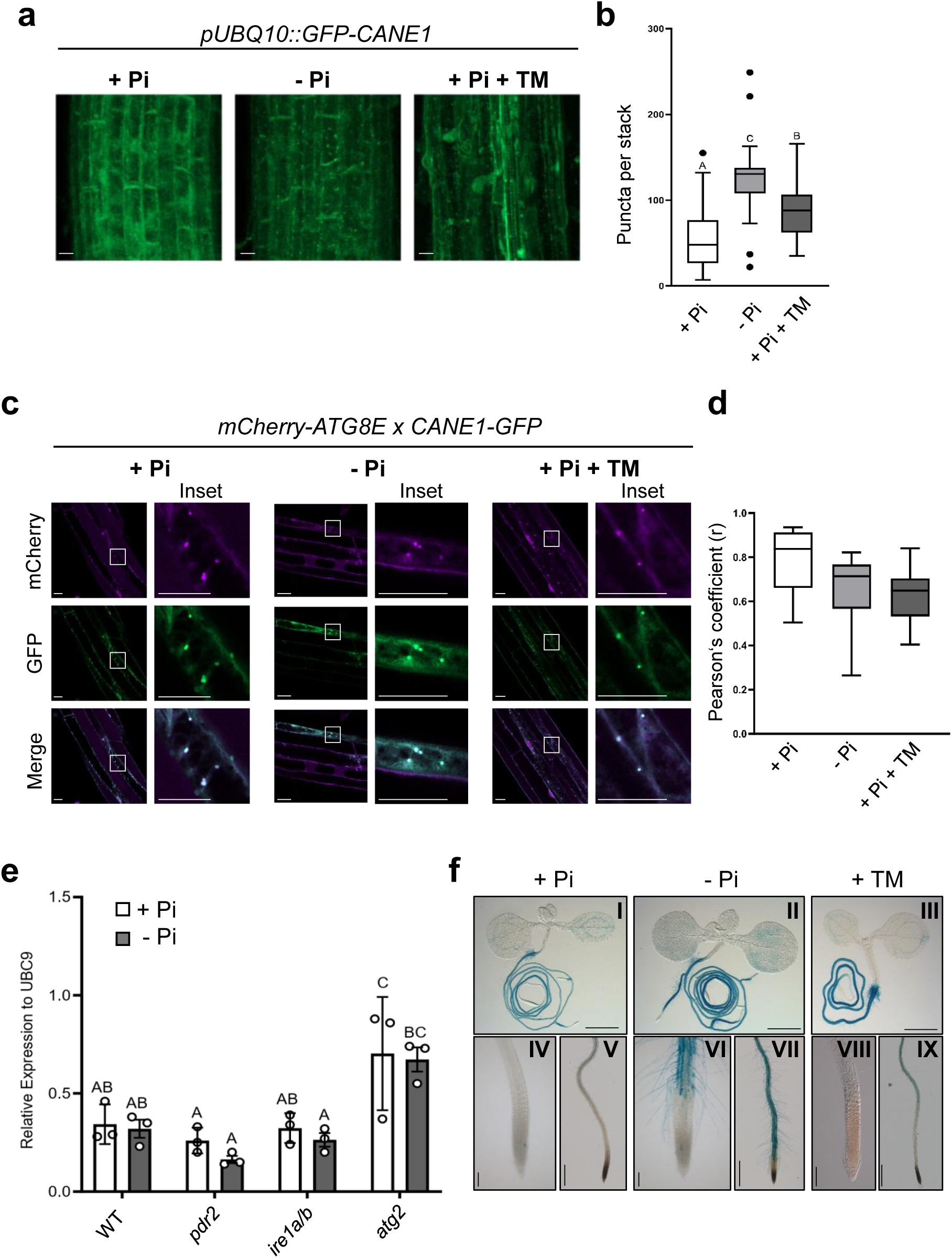
CANE1 protein functions in ER stress-induced autophagy by local Pi sensing. **a**, GFP-CANE1 derived fluorescence in primary root tips of transgenic (*pUBQ10::GFP-CANE1*) wildtype seedlings after germination on +Pi agar (5 d) and subsequent transfer to –Pi medium for 0-24 h or treatment with 5 µg/mL TM for 6 h. Shown are representative images from three independent experiments fused Z-stacks of the transition zone. Scale bar 20 µm. **b**, Quantification of GFP-CANE1 puncta per stack in the indicated treatments. Box plots show medians and interquartile ranges of detected puncta; outliers (>1.53 interquartile range) are shown as black dots. Letters denote statistical differences in condition at p < 0.05 (Two-way ANOVA and Tukey’s HSD post hoc test) and n ≥ 30. **c**, CANE1-GFP puncta colocalize with mCherry-ATG8E *in vivo*. Co-localization analyses of single plane confocal images obtained from transgenic Arabidopsis roots co-expressing mCherry-ATG8E (magenta) with CANE1-GFP (green) in wildtype. Seedlings were germinated 5 days prior transfer to the indicated treatment for 1 day (+/-Pi) or 5 h (TM). Scale bars, 20 µm, n ≥ 10. **d**, Pearson’s Coefficient (r) analysis of the colocalization of CANE1-GFP and mCherry-ATG8E. Box plots show medians and interquartile ranges with five independent measures per biological replicate, n ≥ 10. **e**, *CANE1* expression relative to UBC9 in gain of primary root tip 1 day after transfer of wildtype, *pdr2*, *ire1a/b* and *atg2.* Letters denote statistical differences in condition at p < 0.05 (Two-way ANOVA and Tukey’s HSD post hoc test) and n = 3. **f**, *pCANE1::GUS* activity in 10 days old wildtype seedlings after transfer to the indicated media for two days or incubation with TM for 6 h, n = 3. Overview of the seedlings (I-III) scale bar 1 cm, primary root tip (IV-IX) with IV, VI, VIII scale bar 100 µm and V, VII, IX scale bar 1mm; n ≥ 6.

**Supplementary Figure S8:**
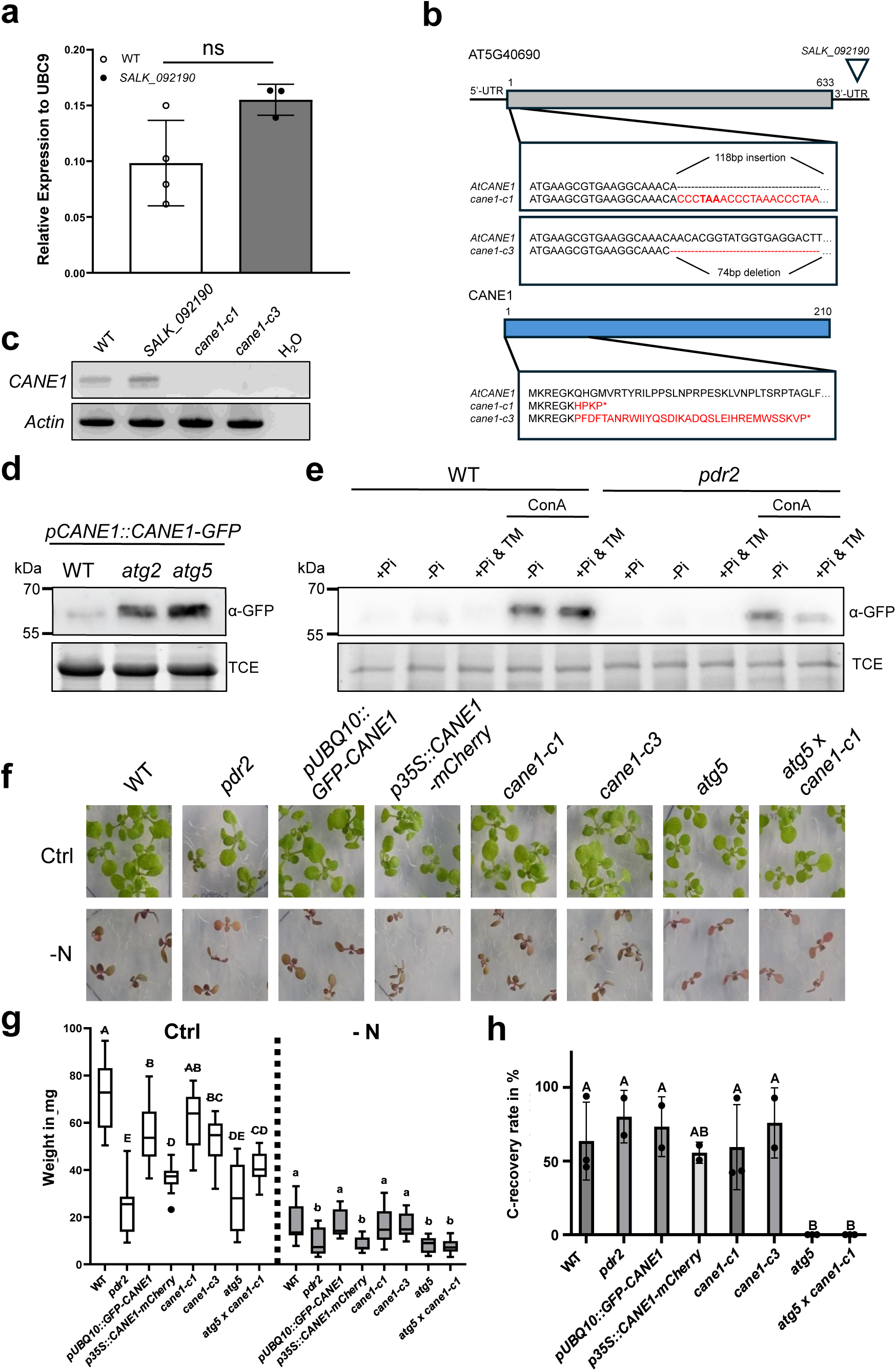
CANE1 functions in ER stress-mediated autophagy. **a,** Transcriptional regulation of *CANE1* is not altered in SALK-Insertion line. Analysis of *CANE1* relative transcript level (normalized to UBC9 expression) of wildtype and *SALK_092190* insertion line seedlings (germinated for 5 days on +Pi media) ±SD; n ≥ 3; Students t-test. **b**, sequence analysis of *CANE1* (AT5G40690). Scheme of two *cane1* CRISPR mutants, *cane1-c1* and *cane1-c3*. Both mutants contain deletions/insertions where *cane1-c1* harbors a 118-bp insertion, while the *cane1-c3* mutant possesses a 74-bp deletion. The deletions/insertions lead to amino acids changes and premature stop codons in both mutants. **c**, Expression analysis of *CANE1* by semiquantitative RT-PCR by using cDNA isolated from wildtype, *SALK_092190*, *cane1-c1*, and *cane1-c3* seedlings. **d**, Amount of *pCANE1::CANE1-GFP* in wildtype, *atg2* and *atg5* seedlings (germinated for 5 days on +Pi media) were analyzed by Western Blot. TCE was used as loading control, n = 4. **e**, 5-day old seedlings expressing *pCANE1::CANE1-GFP* in wildtype and *pdr2* mutant were transferred to the indicated media for 24 h, TM (300 ng/µl) ConA (1µM). Whole roots were harvested and analyzed by immunoblot. TCE was used as loading control, n = 3. **f, g** 5 days-old seedlings of the indicated genotypes were grown on full (Ctrl) and N-depleted media for 7 days. **g**, Analysis of seedling weight after treatment. Box plots show medians and interquartile ranges of primary root gain. Letters denote statistical differences in treatment condition at p < 0.05 (One-way ANOVA and Tukey’s HSD post hoc test) and n = 8-16. **h**, Loss-of *CANE1* does not rescue carbon starvation in *atg5* mutant plants. 5 days-old seedlings of the indicated genotypes were germinated on full media, prior to carbon starvation for 12 days, followed by recovery on full media for one week, ±SD; n = 3; One-way ANOVA, Post-hoc Tuckey, *p value ≤ 0.05.

**Supplementary Figure S9:**
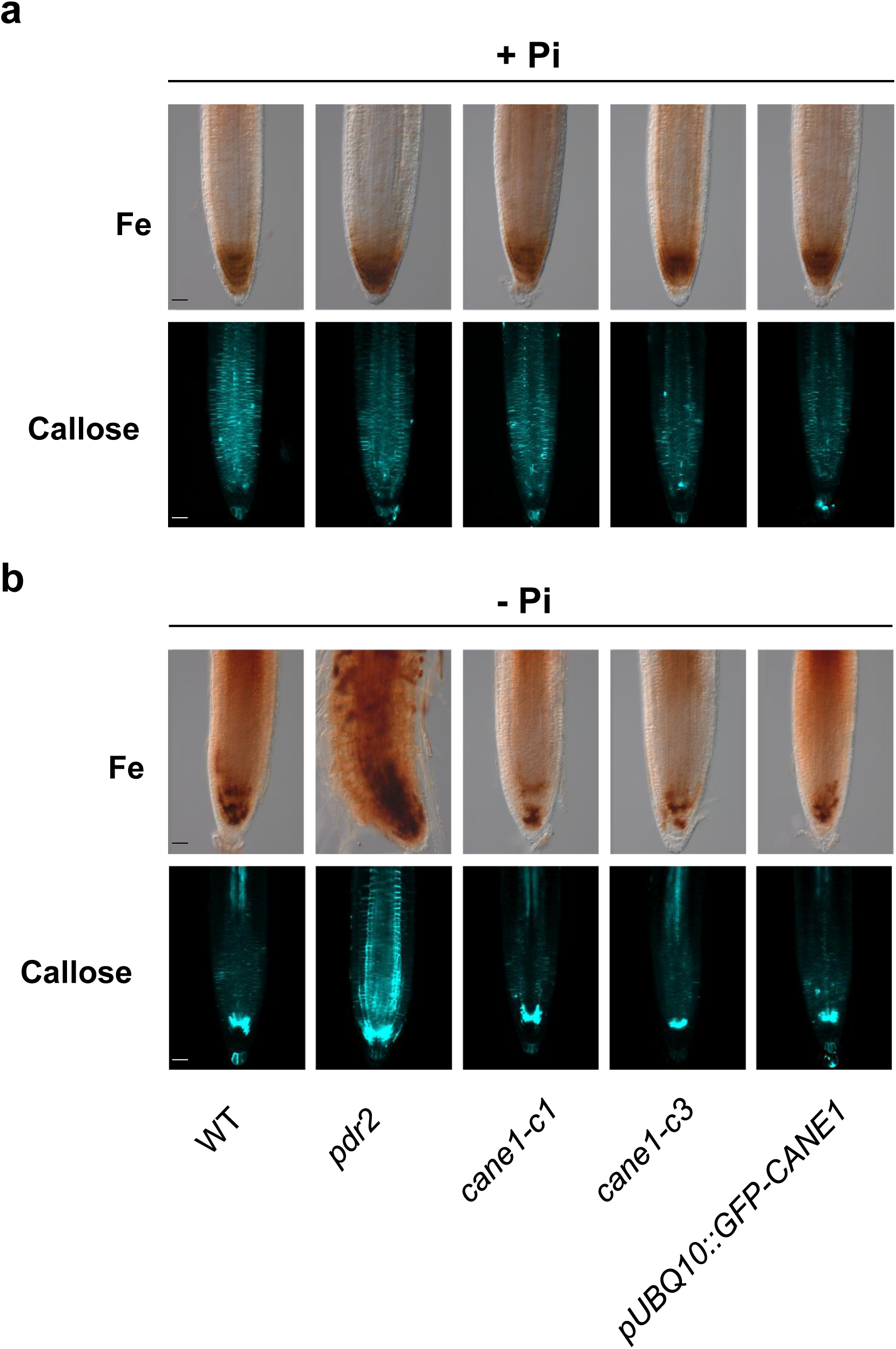
LPR1-mediated local Pi response is not influenced by CANE1. a,b. After germination (5 d, +Pi), seedlings were transferred to +Pi (**a**) or –Pi (**b**) medium (25 μM Fe). Top row: Three days after transfer, root tips of wildtype, *pdr2*, *cane1-c1*, *cane1-c3 and pUBQ10::GFP-CANE1* were monitored for Fe^3+^ accumulation by Perls staining coupled to diaminobenzidine (DAB) intensification. Bottom row: One day after transfer, root tips were monitored for callose deposition by aniline blue staining. Shown are representative images (n ≥ 20). Scale bars 50 μm.

**Supplementary Figure S10:**
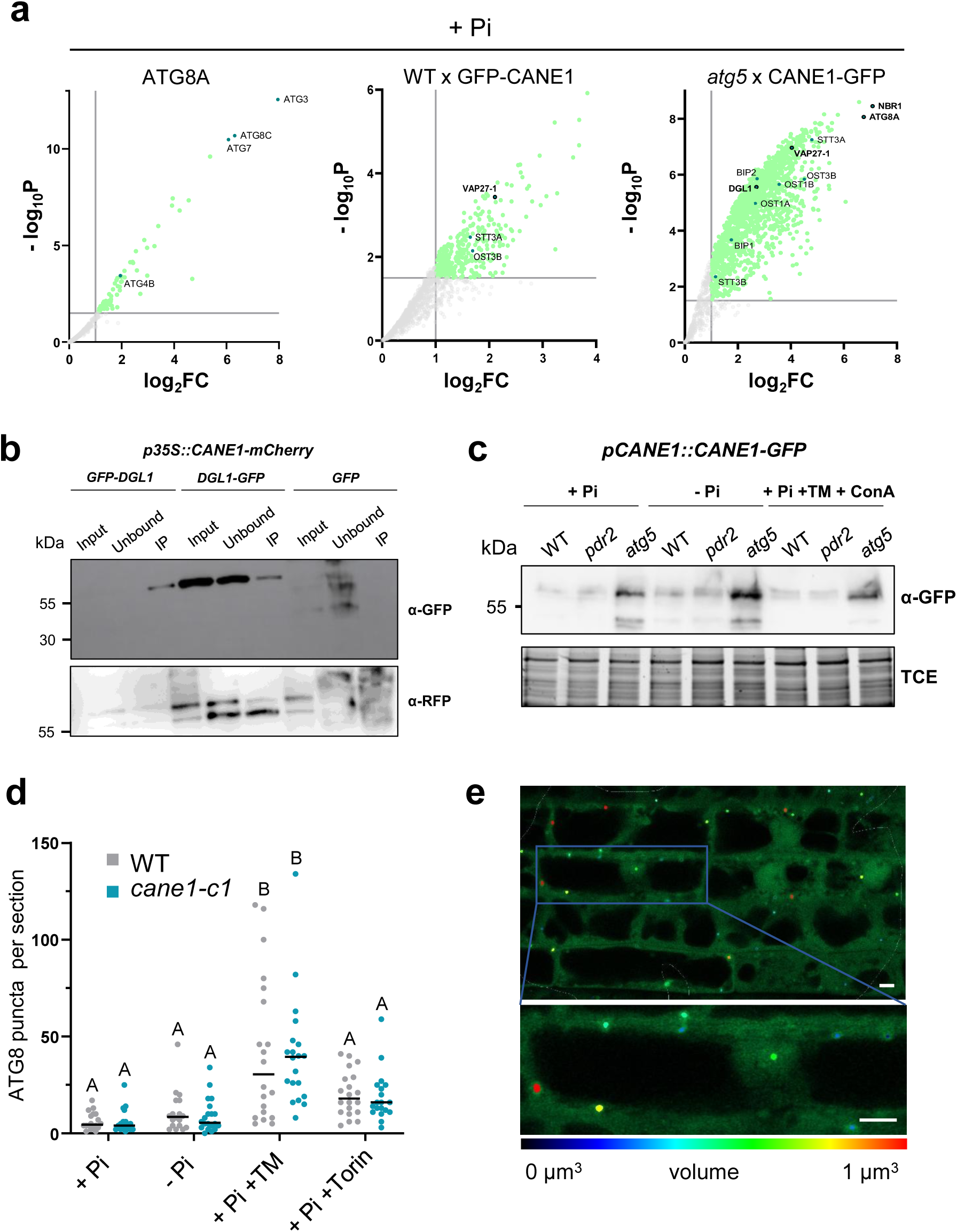
CANE1 intersects with the autophagic pathway in ER stress. **a**, Enrichment of proteins interacting with *p35S::GFP-ATG8A*, *pUBQ10::GFP-CANE1* in wildtype and *pCANE1::CANE1-GFP* in *atg5* background in + Pi compared with a control expressing free GFP and represented by a volcano plot. The horizontal grey line indicates the threshold above which proteins are enriched (P value <0.03, quasi-likelihood negative binomial generalized log-linear model), and the vertical dashed line indicates the threshold where the proteins’ log_2_FC is > 1. **b**, Western blot analysis from immunoprecipitation (IP) experiments of tobacco leaves co-infiltrated with *p35S::CANE1–mCherry* together with *p35S::DGL1-GFP*, *pUBQ10::GFP-DGL1* or *p35S::GFP-HDEL.* Input and bound proteins were visualized with anti-GFP and anti-RFP antibodies, n = 3. **c**, Amount of *pCANE1::CANE1-GFP* in wildtype, *pdr2* and *atg5* roots were analyzed by immunoblot. Seedlings were germinated 5 days on +Pi prior to transfer to the indicated media for 24 h or ER stress treatment with 5 µg/ml TM and 1 µM ConA for 6 h. TCE was used as loading control, n = 4. **d**, Autophagosome quantification in roots of wildtype and *cane1-c1* expressing *p35S::GFP-ATG8A.* Seedlings were germinated for 5 days on +Pi media prior to transfer for 24 h to +Pi or –Pi media or incubated with 5 µg/ml TM or 1µM torin for 6 h for autophagic induction. Scatter plot, median is shown as bold line, letters denote statistical differences at p < 0.05 (Two-way ANOVA and Tukey’s HSD post hoc test), n ≥ 18. **e**, Representative image of autophagosome volume quantification in Arivis. *p35S::GFP-ATG8A in* wildtype 24 h post transfer to –Pi is shown. Scale = 5 µm. Color-coded autophagosome volume according to the scale.

